# Transmissible cancers and the evolution of sex under the Red Queen hypothesis

**DOI:** 10.1101/2020.04.01.019885

**Authors:** Thomas G. Aubier, Matthias Galipaud, E. Yagmur Erten, Hanna Kokko

## Abstract

The predominance of sexual reproduction in eukaryotes remains paradoxical in evolutionary theory. Of the hypotheses proposed to resolve this paradox, the “Red Queen hypothesis” emphasizes the potential of antagonistic interactions to cause fluctuating selection, which favours the evolution and maintenance of sex. While empirical and theoretical developments have focused on host-parasite interactions, the premises of the Red Queen theory apply equally well to any type of antagonistic interactions. Recently, it has been suggested that early multicellular organisms with basic anticancer defenses were presumably plagued by antagonistic interactions with transmissible cancers, and that this could have played a pivotal role in the evolution of sex. Here, we dissect this argument using a population genetic model. One fundamental aspect distinguishing transmissible cancers from other parasites is the continual production of cancerous cell lines from hosts’ own tissues. We show that this influx dampens fluctuating selection and therefore makes the evolution of sex more difficult than in standard Red Queen models. Although coevolutionary cycling can remain sufficient to select for sex under some parameter regions of our model, we show that the size of those regions shrinks once we account for epidemiological constraints. Altogether, our results suggest that horizontal transmission of cancerous cells is unlikely to cause fluctuating selection favouring sexual reproduction. Nonetheless, we confirm that vertical transmission of cancerous cells can promote the evolution of sex through a separate mechanism, known as similarity selection, that does not depend on coevolutionary fluctuations.

## INTRODUCTION

Sexual reproduction entails several and often severe costs (Lehtonen et al., 2012), yet most eukaryotes engage in sex, at least occasionally (Speijer et al., 2015). To explain this apparent paradox, much theory has been developed to identify the benefits associated with sexual reproduction (Otto and Lenormand, 2002; Agrawal, 2006a; Hartfield and Keightley, 2012; Schwander et al., 2014). Sex shuffles genetic material from parent individuals and breaks apart allele combinations built by past selection. Whether this is selected for depends strongly on the stability of the environment. In a stable environment, selection is likely to have already brought favourable combinations of alleles together in the past, and continuing genetic mixing can become deleterious (Feldman, 1972; Barton, 1995). In many models, therefore, the evolution of sex relies on the advantage that lineages receive from mixing genetic materials in an environment that is not stable (Otto and Michalakis, 1998). Under a scenario of fluctuating selection, sex can be beneficial because it breaks apart allele combinations that have been built by past selection and that by now have become disadvantageous (Barton, 1995; Gandon and Otto, 2007).

The “Red Queen hypothesis” for the evolution of sex emphasizes the potential of host-parasite interactions to cause fluctuating selection, thus favouring genetic mixing (Haldane, 1949; Hamilton, 1975; Levin, 1975; Hamilton, 1980; Bell, 1982; Lively, 2010; Gibson et al., 2018) (not to be confused with the macroevolutionary Red Queen hypothesis; Van Valen, 1973). Hosts with a rare genotype are often less susceptible to common parasite strains (e.g., Lively and Dybdahl, 2000), and the resulting coevolution between hosts and parasites (so-called ‘Red Queen dynamics’) yields negative frequencydependent selection, such that selection fluctuates over time for host and parasite alike (Bell and Maynard Smith, 1987). Genetic mixing is then favoured because it can produce offspring with genetic associations that are currently rare and therefore less susceptible to parasites (Hamilton, 1980; Bell and Maynard Smith, 1987; Nee, 1989).

Until now, studies of the Red Queen hypothesis have considered hosts and parasites that belong to distinct taxonomic groups. In a recent opinion piece, Thomas et al. (2019) break from this tradition and highlight the intriguing potential role of transmissible cancerous cell lineages for the evolution of sex. Cancer manifests itself as somatic cells breaking free of multicellular cooperation and proliferating uncontrollably, often at the cost of the organism’s (as well as the cancer cells’) life (Aktipis et al., 2015). Cancer is an evolutionary dead end (Sidow and Spies, 2015), but an exception arises when cancer cells are transmissible and outlive their host, behaving in this respect identically to parasites that infect new hosts. In this latter case, hosts can suffer not only from cancer cells arising from their own tissues but also from transmitted cancer cells that originated in other host individuals.

Transmissible cancers are fundamentally different from contagious agents that elevate cancer risk (e.g., human papillomavirus causing cervical cancers; Bouvard et al., 2009). Transmissible cancers are directly transmitted to new hosts, and do not require cells or viral particles of another taxon to play any role in the infection system. So far, transmissible cancerous cell lines have been observed in a few taxa only, namely three mammal species (Ashbel, 1945; Murgia et al., 2006; Murchison, 2008; Pye et al., 2016) and four bivalve species (Metzger et al., 2015, 2016; Yonemitsu et al., 2019), but early multicellular organisms, with presumably basic anticancer defenses, may have been plagued by this problem more than extant ones (as argued by Ujvari et al., 2016b; Thomas et al., 2019).

On this basis, Thomas et al. (2019) propose the intriguing hypothesis that the prevalence of sex in multicellular eukaryotes may have been originally driven by transmissible cancerous cell lines regularly infecting multicellular hosts (note however that eukaryotes tend to be sexual even if they are unicellular). According to this view, the diversity of genotypes created by sex helps individuals in the task of differentiating between self and non-self, thus reducing susceptibility to transmissible cancers. Selection on the multicellular host to avoid infection by transmissible cancer is therefore akin to selection induced by heterospecific parasitic agents, favouring the evolution and maintenance of sex.

Transmissible cancers are indeed similar to other parasites in that the long-term survival of their lineages depends on their successful transmission to other hosts, which requires circumventing the host’s immune system. Transmissible cancer cells, however, present a particularly thorny problem for the host as their genetic makeup (and hence the cellular phenotype) is by default very similar to that of the host, since they originate via mutation from the host’s own tissue or from a conspecific tissue (but see also Metzger et al., 2016, for a case of cross-species transmission). Self/non-self recognition is conceptually similar to the ‘matching-alleles’ model of host defence against parasites (relevant for the Red Queen process; e.g., Bell and Maynard Smith, 1987; Seger and Hamilton, 1988; Howard and Lively, 1994; Agrawal and Lively, 2001, 2002), in that the cancer cell’s infection prospects depend on the genotypic composition of the host population. At first glance, this sets the scene for antagonistic coevolution between the transmissible cancer and its host, favouring the evolution of sex just as predicted under the Red Queen hypothesis. Without formal inquiries, however, it is difficult to judge whether antagonistic interactions between hosts and transmissible cancers lead to fluctuating selection of the type that is essential for Red Queen dynamics to take place. Specifically, it is unclear whether fluctuating selection can be maintained when susceptible hosts themselves produce the parasitic cancerous lines to which they are susceptible.

In this paper, we investigate whether antagonistic interactions between hosts and transmissible cancers can promote the evolution of sex under the Red Queen hypothesis. We analyze a population genetic model of fluctuating selection and complement it with an epidemiological model. The latter model builds an explicit epidemiological setting that we then use to examine the likely parameter values that the population genetic model takes. This combined use of two models allows us to evaluate how likely it is for the modelled system to find itself within a selection regime where Red Queen dynamics can favour sexual reproduction.

Our approach is an intentionally simplified version of all self/non-self recognition systems (with only two loci involved in recognition, plus one modifier that determines sexual/asexual reproduction), and we regard it as the first necessary step for understanding the conditions under which interactions between hosts and transmissible cancers can yield to Red Queen dynamics promoting the evolution of sex. We focus on one of the fundamental aspects that distinguish transmissible cancers from other parasites: the continual production of cancerous cell lines from hosts’ tissues (a process that we refer to as ‘neoplasia’), which we show to inhibit the evolution of sex under the Red Queen hypothesis. The inhibitory effect arises because ‘neocancers’ produced via neoplasia are likely to infect hosts with a common genotype, strongly reducing the lag between hosts and transmissible cancers evolutionary dynamics, which is necessary for coevolutionary fluctuations to occur. Coevolutionary dynamics can select for sex under some parameter regions of the population genetic model, yet we show that the size of those regions shrinks once we account for epidemiological constraints.

## 1. POPULATION GENETICS

### 1.1. Population genetic model

We extend a standard population genetic model of the Red Queen Hypothesis (e.g., Otto and Nuismer, 2004; Salathé et al., 2008, 2009; Engelstädter and Bonhoeffer, 2009) to account for neoplasia, i.e. the fact that cancers originate from conspecific hosts and bring their genotypes into the population of transmissible cancer cells. We distinguish between two stages that characterize transmissible cancer cells: cancer cells successfully transmitted to a new host in a previous generation (called ‘transmitted cancers’), and those that do not yet have such an infection history but are directly derived from the original host where neoplasia occurred (called ‘neocancers’). Neocancers become transmitted cancers as soon as they successfully infect a new host. We specifically test whether coevolution between hosts and transmissible cancers can favour the evolution of sex under the Red Queen hypothesis.

We follow the genotypic frequencies of haploid hosts and haploid cancer cells through a life cycle that consists of a census, reproduction, neoplasia (development of neocancers), and selection (that depends on interactions between hosts and transmissible cancers). We assume that hosts and cancer cells each form populations of sufficient size such that we can ignore the effects of genetic drift. We also assume that time is discrete.

#### Genotypes

Hosts and transmissible cancers are haploid and have two loci, *A* and *B*, with two possible alleles (A/a, B/b) that determine the outcome of the interaction between hosts and cancers.

Hosts possess an additional modifier locus *M* with two possible alleles (M/m) that determine whether the host reproduces sexually or asexually (here, gene order of *A*, *B* and *M* does not matter). Hosts carrying the allele M reproduce sexually, whereas hosts carrying the allele m reproduce asexually. Although cancer cells are ultimately derived from host cells and thus carry the modifier locus *M* too, we assume for simplicity that they never engage in sex and recombination, consequently there is no need to distinguish between cancer cells with alleles m and M in the model. Our approach does not give cancer cells themselves the ability to fuse and recombine (Miroshnychenko et al., 2020; Pienta et al., 2020), as this allows us to follow the original argumentation of Thomas et al. (2019).

As a whole, therefore, hosts are of eight possible genotypes (mAB, mAb, maB, mab, MAB, MAb, MaB, Mab) whose frequencies are 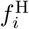, and transmissible cancer cells are of four possible genotypes (AB, Ab, aB, ab) whose frequencies are 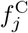.

#### Reproduction

Host individuals carrying the allele M at the modifier locus *M* engage in sexual reproduction. Random mating (which brings two haploid genomes together) is followed immediately by meiosis. During meiosis, loci *A* and *B* recombine at rate *r*_host_. Recombination between these loci (*A* and *B*), and locus *M* is irrelevant, since sexual progeny inherit the allele *M* from both parents. By contrast, host individuals carrying allele m at locus *M* engage in asexual reproduction, and genotypic frequencies in the asexual host population do not change. All transmissible cancer cells reproduce clonally, i.e., genotypic frequencies in transmissible cancer cells do not change.

Mutations at the interaction loci occur at a rate of *m*_host_ and *m*_cancer_ per locus per generation in the hosts and the cancer cells, respectively. If a mutation occurs, allele A (resp. B) becomes allele a (resp. b), and vice versa. We assume there is no mutation at the modifier locus *M*, since our aim is to assess whether an allele M controlling sexual reproduction can invade the population once it is introduced in the population.

Overall, from frequencies 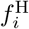 and 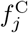, we can calculate the genotypic frequencies 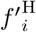 and 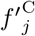 in hosts and cancer cells after reproduction, assuming that generations do not overlap.

#### Host neoplasia and development of neocancers

Transmissible cancers are ultimately derived from neoplasia, and have initially the same genotype as the neocancer’s original host (i.e., we assume no somatic mutation/selection during the oncogenic process that produces the cancer within the original host). We use *α* to denote the proportion of transmissible cancers that are ‘neocancers’ (without infection history), with genotypic frequencies 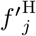 at the interaction loci. The remaining proportion 1 − *α* of cancers are ‘transmitted cancers’ (with infection history), with genotypic frequencies 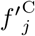 differing from those of neocancers. If *α* = 0, neoplasia does not occur, and transmissible cancers are in that case best seen as classical heterospecific parasites that do not arise from host cells themselves.

Therefore, after neoplasia (the development of neocancers) has occurred, genotypic frequencies in the transmissible cancers are:

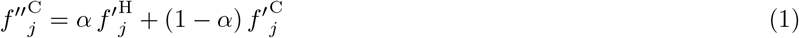

We assume that neoplasia does not depend on host genotype – i.e., hosts of any genotype develop neocancers at the same rate. We also assume that host genotype does not impact the severity of fitness consequences (to the host) of neoplasia. Therefore, even if neoplasia increases host mortality, this does not change genotypic frequencies in the host population: 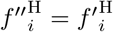.

#### Selection

During the selection phase, hosts are assumed to encounter transmissible cancers (neocancers and transmitted cancers) proportionally to their frequency. Changes in genotypic frequencies are determined by the match/mismatch at the interaction loci (*A* and *B*).

We implement the commonly used matching-alleles interaction model, now interpreted in the context of self/non-self recognition. When there is an exact match between genotypes *i* and *j* of the host and the infecting cancer, the infecting cancer has high fitness (which we model as fitness coefficient 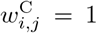), while the host suffers a fitness cost (fitness coefficient 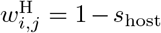). In the alternative case where the genotypes *i* and *j* of the host and the infecting cancer do not match, the host retains its high fitness advantage (fitness coefficient 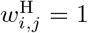), while the infecting cancer suffers a fitness cost (fitness coefficient 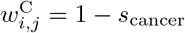).

Assuming that the probability of interaction between host genotype *i* and cancer genotype *j* is the product of their respective frequencies, 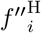 and 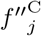, the frequency of host genotype *i* after selection is 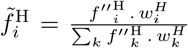, where 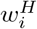 is the fitness of host genotype *j*, given by 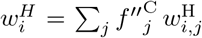. The genotypic frequencies in cancer after selection, 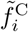, are calculated analogously (with fitness coefficients 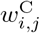).

During the selection process presented above, we therefore made the assumption that transmission occurs only horizontally. In Appendix B, however, we implement vertical transmission of cancerous cells from mother to offspring. Vertical transmission causes ‘similarity selection’ that has also the potential to favour sex and recombination, while being distinct from genotypic selection created by coevolutionary fluctuations in standard Red Queen models (Agrawal, 2006b). Agrawal (2006b) used a standard population genetic approach to emphasize the importance of similarity selection for the evolution of sex, but more theoretical and empirical work is required to confirm that this is a potent mechanism favouring sexual reproduction. To our knowledge, the only other models considering similarity selection in other evolutionary contexts are those of Greenspoon and M’Gonigle (2013, 2014).

#### Numerical simulations

We initiate the populations assuming that all hosts are asexual (in hosts, the frequency of allele m at locus *M* is set to 1), and other allele frequencies are initialized randomly; i.e., genotypic frequencies are drawn following uniform distributions over the range [0, 1], and are then normalized such that their sum is equal to 1 in hosts and transmissible cancers.

The initial host and cancer populations are allowed to coevolve for 10, 000 generations (burn-in period), with the dynamics computed using the recursion equations above. The mutant allele M is then introduced such that 5% of the host population becomes sexual (switches from allele m to allele M), and host and cancer populations are allowed to coevolve for 1, 000, 000 generations. For each combination of parameters tested, we perform 100 simulations characterized by different initial conditions, such that coevolutionary dynamics can explore different limit cycling dynamics. If we find the frequency of allele M to reach and maintain a frequency > 0.999 for at least 500 generations over the course of at least one simulation, we assume that sex can invade under the combination of parameters implemented.

To provide a sensitivity analysis, we vary the parameters *α* (proportion of neocancers among the population of transmissible cancers) and (*s*_host_, *s*_cancer_) (selection coefficient associated with the interaction between hosts and cancers), with mutation rates *m*_host_ = *m*_cancer_ = 10^−5^. For each combination of parameters, we run simulations with different values of *r*_host_ ∈ {0.005, 0.01, 0.02, 0.05, 0.1, 0.2, 0.3, 0.4, 0.5} (recombination rate if hosts reproduce sexually). If sex can invade for at least one value of *r*_host_, then we consider that sex can evolve under the combination of parameters (*α, s*_host_, *s*_cancer_) tested. Note that our assumption of implementing *α* as a parameter is a simplification that we make to aid conceptual understanding of the role that newly arisen cancers play in Red Queen dynamics; we thereafter switch to viewing *α* as an emergent property of the system in the epidemiological model explained in section 2, below.

### 1.2. Red Queen dynamics and evolution of sex

Under the Red Queen hypothesis, antagonistic interactions between hosts and parasites cause fluctuating selection leading to non-steady coevolutionary dynamics (Red Queen dynamics) that favours the evolution of sex. Without neoplasia (*α* = 0), we show that such dynamics occur in the our three-locus population genetic model (cf. parameter spaces delimited by the plain red lines in Fig. 1A; e.g., Fig. 1B). Consequently, sexual reproduction can invade in the host population (in purple in Fig. 1A), especially if it associates with an intermediate recombination rate (Fig. S1). These predictions are in agreement with previous theory on the Red Queen hypothesis.

**Figure 1:**
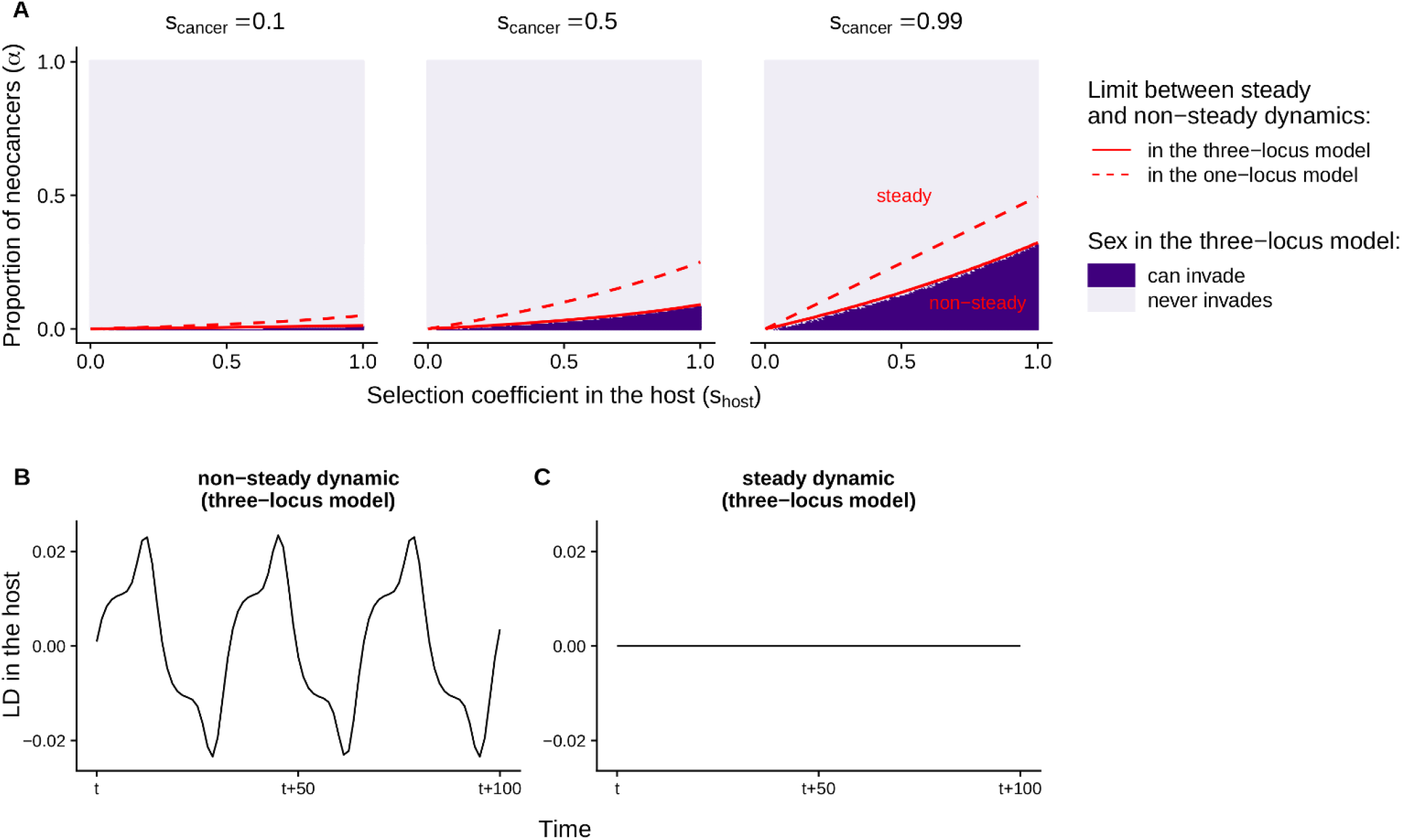
Coevolutionary dynamics between hosts and transmissible cancers, and evolution of sex. A. Sensitivity of the population genetic models to the selection coefficients (*s*_host_, *s*_cancer_) and to the proportion of transmissible neocancers that are recently derived from the original host (*α*). Red lines delimit the parameter spaces leading to non-steady and steady coevolutionary dynamics in the three-locus model (plain line; found numerically) and in the simplified one-locus model (dashed line; found analytically in Appendix A). In the three-locus model, the dynamic is defined as ‘steady’ when the variance in genotypic frequencies over 500 time steps is below 10^−10^. Dark purple indicates conditions under which a modifier allele associated with sexual reproduction (and with recombination, at least for one of the recombination rates tested) can invade in the three-locus model in at least one of the 100 simulation runs. B-C. Examples of non-steady and steady coevolutionary dynamics in the three-locus model. The linkage disequilibrium in the host is calculated as 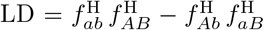, i.e., a positive linkage disequilibrium here represents a non-random excess of homozygotes. In subfigures B and C, parameter values are: *s*_host_ = 0.5, *s*_cancer_ = 0.8, and *α* = 0 (B) or *α* = 0.1 (C).

The results change when some of the transmissible cancers in circulation are neocancers, the results of neoplasia occurring in the original hosts (*α* > 0). Even a small proportion of neocancers is sufficient to bring the coevolutionary dynamics between hosts and transmissible cancers to a halt (Fig. 1A; e.g., Fig. 1C). Neoplasia tightens the link between genotypic frequencies in host and cancer (Equation 1), in the sense of reducing any time lag between the two evolutionary dynamics. Since a time lag is necessary for coevolutionary fluctuations to occur and for sex to be favoured in Red Queen models, the continual production of new cancer cells from original hosts makes it impossible for sexual reproduction to invade if the proportion of neocancers is too high (Fig. 1A).

Other strong determinants of the coevolutionary dynamics are the strengths of selection associated with the interaction between hosts and transmissible cancers (*s*_host_, *s*_cancer_). Overall, increased strengths of selection promote non-steady coevolutionary dynamics and favour the evolution of sex (Fig. 1A). In other words, when resistance or infection associate with a high fitness in host and cancer respectively, Red Queen dynamics can occur and favours the evolution of sex.

Notably we get the same results when we consider that the ancestral reproductive mode is facultative sexual reproduction (with hosts originally reproducing sexually and asexually to the same extent; Fig. S2). Neoplasia also dampens coevolutionary cycling when we consider more than two interaction loci mediating the interaction between hosts and cancers (Fig. S3); note that increasing the genotypic space constrains the conditions under which sex can evolve under the Red Queen hypothesis (as shown by Otto and Nuismer, 2004).

To gain further insights into the effect of the proportion of neocancers (*α*) on coevolutionary dynamics between hosts and transmissible cancers, we consider a simplified one-locus population genetic model and solve it analytically. This model is based on a single autosomal haploid locus *A* with two possible alleles (A/a), controlling the interaction between hosts and transmissible cancers. This model does not include a modifier locus controlling the reproduction mode of the host; all hosts are reproducing asexually. We also assume that there is no mutation (*m*_host_ = *m*_cancer_ = 0). We determine the local stability of all equilibria by analyzing the eigenvalues of the corresponding Jacobian matrices (Appendix A). We show that all equilibria are unstable, leading to non-steady coevolutionary dynamics, only if the proportion of neocancers is lower than a threshold value *A*\*(*s*_host_, *s*_cancer_):

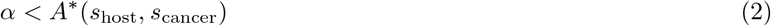

With:

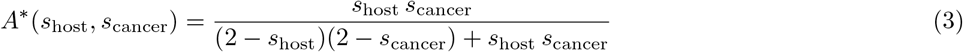

This condition is represented by a dashed red line in Figure 1A. The maximum value of *α* that allows for a non-steady coevolutionary dynamic becomes higher as selection coefficients increase (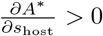 and 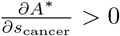), as found numerically in the model with two interaction loci (plain red line in Fig. 1A). This analytical derivation reveals that the amplitude of fluctuations in selection is not merely getting small, but is deterministically shrinking to zero as *α* increases. Therefore, in this simplified genetic setting, neoplasia always dampens the Red Queen dynamics. This explains why in the three-locus model, sex cannot evolve if the proportion of transmissible cancers deriving from host neoplasia is too high.

Thus far we have considered horizontal transmission only, which in our model fails to promote the evolution of sex as soon as neoplasia dampens coevolutionary fluctuations. While the theory on the Red Queen hypothesis relies on non-steady coevolutionary dynamics, antagonistic interactions can favour the evolution of sexual reproduction via other processes. In Appendix B, we show that vertical transmission of cancerous cells can promote the evolution of sex through a separate mechanism, called similarity selection (Agrawal, 2006b), where the sex-promoting effect operates in the absence of coevolutionary fluctuations. Similarity selection occurs when there is a cost to being genotypically similar to one’s family members. In particular, due to the transmission of cancerous cells from parent to offspring, this cost exists if infection compatibility is under genetic influence (e.g., as in the matching alleles system of our population genetic model).

## 2. EPIDEMIOLOGY

### 2.1. Epidemiological model

The population genetic model presented above treats the proportion of transmissible neocancers originating via neoplasia (*α*) and the selection coefficient caused by infection by transmissible cancers (*s*_host_) as independent parameters. Yet, these two parameters are likely to be linked in a more realistic epidemiological context – i.e., with some combinations of (*α*, *s*_host_) more likely than others. We therefore build an explicit epidemiological model to determine the epidemiological settings that may favour the evolution of sex (as predicted by the population genetic model, above).

In our epidemiological model, host individuals can be cancer-free (‘susceptible’), develop a neocancer (through neoplasia), or be infected by a transmitted cancer (see the flowchart of the model, Fig. 2). Notably, infected hosts incur an elevated mortality rate. We do not model the possibility of a host having both types of cancers simultaneously, but include a parameter (*θ*) that allows infection status to change from one type to another. Thus, *θ* controls whether neoplasia can ‘take over’ if the host has a transmitted cancer already and whether, conversely, a host undergoing neoplasia can become infected by another cancer.

**Figure 2:**
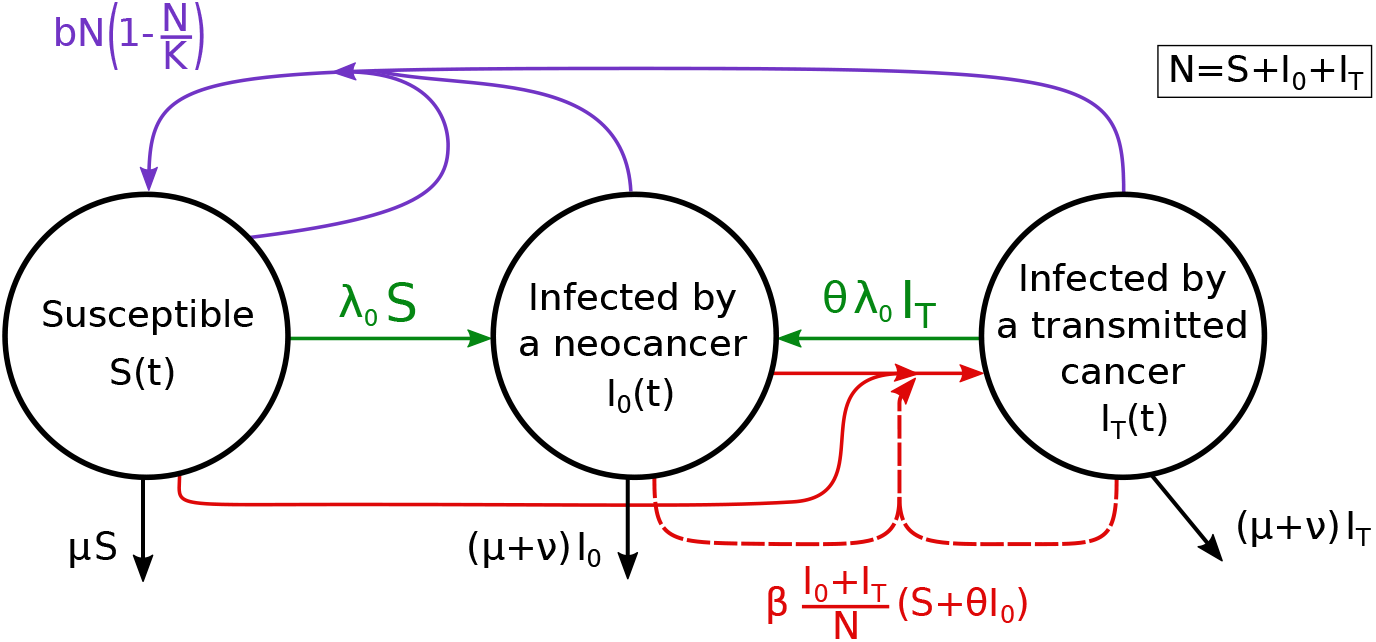
Flowchart of the epidemiological model.

Since the purpose of the model is to investigate likely values of *α* depending on *s*_host_, rather than track any Red Queen dynamics, we do not include any variance in hosts’ and parasites’ genotypes. Instead, we assume that no host can recognize a transmissible cancer as non-self and fend it off; for the sake of analytical tractability, we thus underestimate the proportion of neocancers and overestimate the selection coefficient, by making it easy for cancers to continue infecting hosts beyond the original one (as shown in Appendix D).

#### Ordinary differential equations

The following equations control the changes in densities of susceptible hosts (*S*), hosts that developed a neocancer by neoplasia (*I*_0_), and hosts that are infected by a transmitted cancer (*I*_T_):

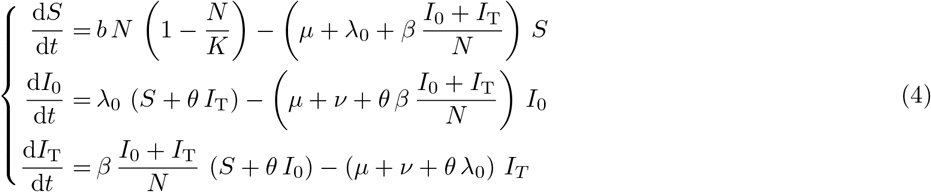

With *N* = *S* + *I*_0_ + *I*_T_.

The host birth rate is density-dependent with a baseline birth rate *b* and a carrying capacity *K*. Baseline mortality rate, independent of density or infection status, is *μ*, which is elevated to *μ* + *ν* in infected hosts (thus *ν* denotes the additional host mortality caused by the cancer). Neoplasia makes hosts develop neocancers at a rate *λ*_0_, and hosts can become infected by a transmitted cancer, controlled by a rate *β* and dependent on the prevalence of transmissible cancers. Parameter *θ* controls the change in infection status. If *θ* = 0, one individual can only ever host one type of cancer. If *θ* > 0, hosts can change infection status.

### 2.2. Equilibrium state

#### Prevalence of transmissible cancers

Depending on the parameter values, the host population either goes extinct or persists with *S*\* + *I*_0_* + *I*_T_* > 0 at equilibrium (Appendix C). This equilibrium state is stable and features an endemic infection by transmissible cancer (*I*_0_* > 0 and *I*_T_* > 0; Appendix C). Note that an equilibrium where all hosts are susceptible, which is possible in standard epidemiological models (SI, SIR and SIS models), is not a feature of our model because susceptible individuals continually produce neocancers.

At equilibrium, we can derive the expression of the prevalence *P* * of transmissible cancers, defined as 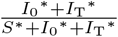 (Appendix C; see also Fig. S4):

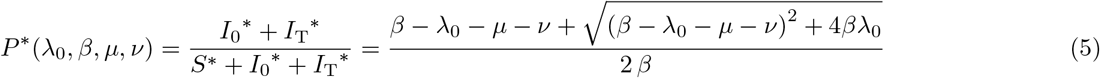

#### Proportion of neocancers

At equilibrium, we determine the expression of the proportion of neocancers 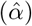 that we define as 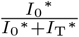 (Appendix C):

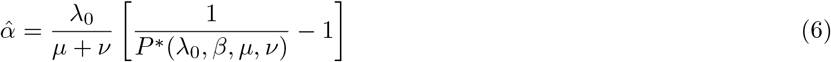

#### Strength of selection due to transmissible cancers

We also determine the expression of the selection coefficient caused by transmissible cancers 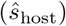 that we define as the mean lifespan reduction due to the risk of being infected by transmissible cancers (Appendix C):

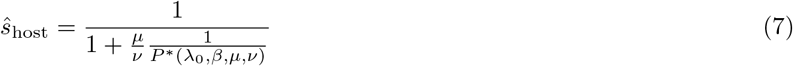

### 2.3. Epidemiological setting favouring the evolution of sex

To infer the epidemiological conditions that can favour the evolution of sex, we compare the properties of the equilibrium state 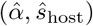 to the conditions that favour the evolution of sex in the previous three-locus population genetic model, assuming the approximations 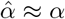 and 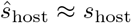.

Some parameters – the birth rate (*b*), the carrying capacity (*K*) and the changes in infection status (*θ*) – have no effect on the proportion of neocancers 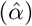 and on the strength of selection caused by transmissible cancers 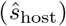 at the equilibrium (those parameters do however affect the dynamics of our epidemiological model before reaching the equilibrium state). By contrast, other parameters – *λ*_0_, *β*, *μ* and *ν* – affect 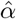 and 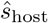 either directly or via their effect on the prevalence of transmissible cancers (*P* *) (Equations 6–7; Fig. 3 and 4).

**Figure 3:**
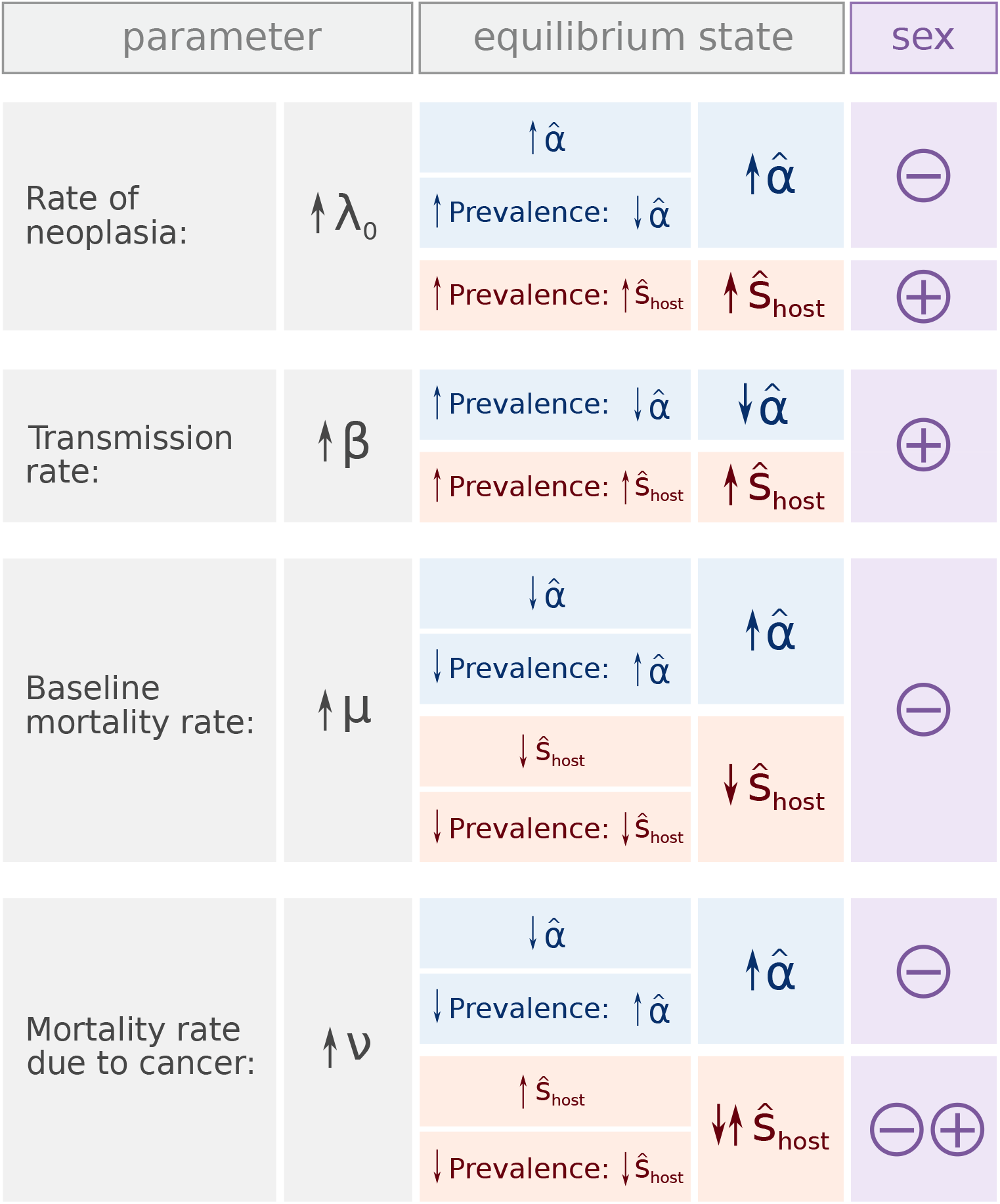
Sensitivity to epidemiological parameters, based on the signs of partial derivatives (see Appendix C). Each epidemiological parameter can affect the proportion of neocancers (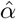; in blue) and the strength of selection caused by transmissible cancer (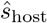; in red) at equilibrium, either directly or via its effect on prevalence of transmissible cancers (cf. Equations 6–7). The right panel under the column “equilibrium state” denotes the overall effect of a change in parameter value on 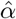 or 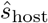. From these sensitivity analyses, we infer whether changes in the epidemiological settings can favour the evolution of sex or not (as predicted in the previous population genetic model; in purple). Note that 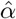 and 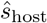 do not depend on parameters *b*, *K*, and *θ*.

**Figure 4:**
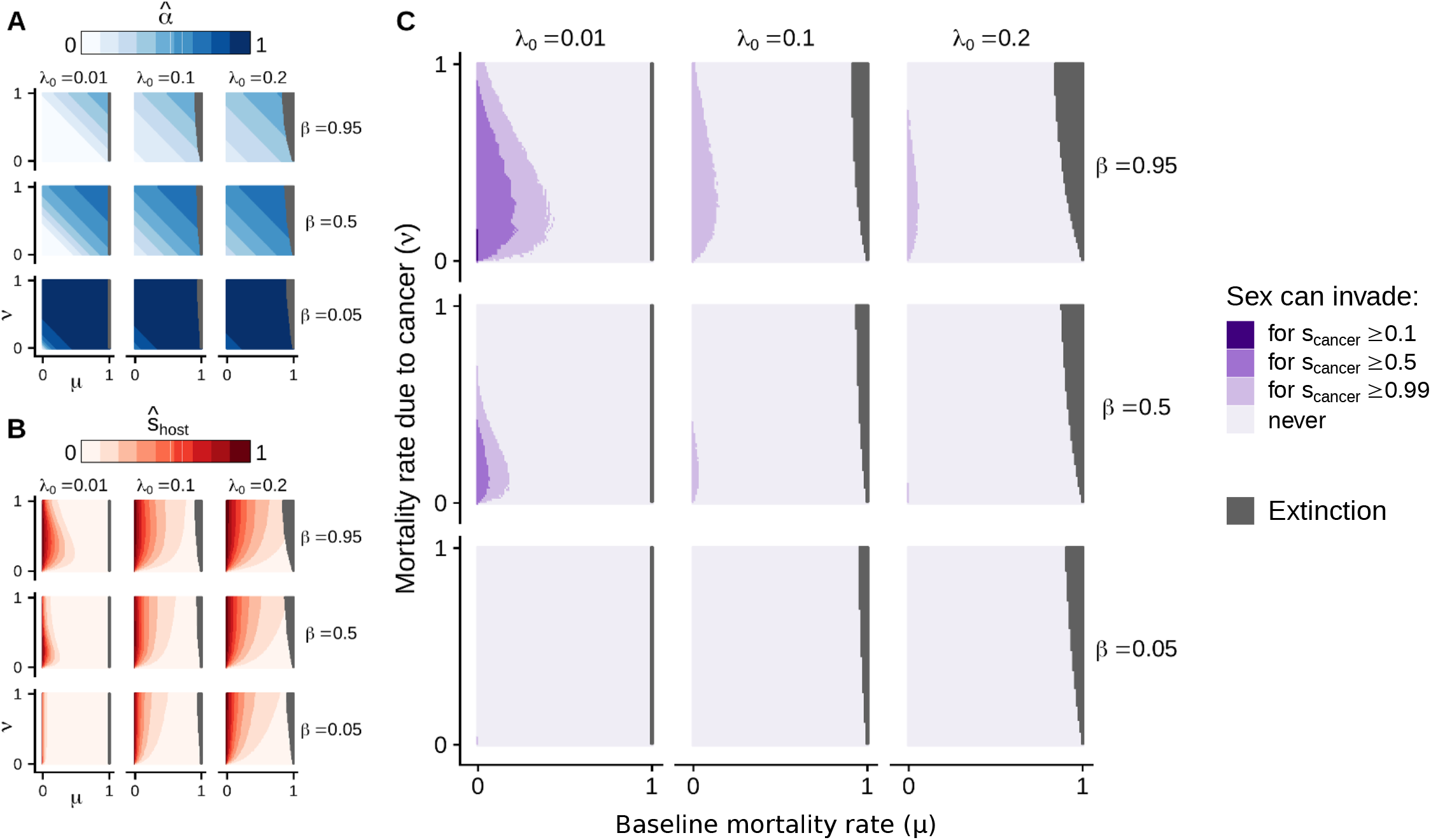
Effects of epidemiological parameters on 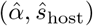 at equilibrium (A, B), and conditions under which sex should be favoured (inferred from the three-locus population genetic model; Fig. 1A) (C). In gray, we represent the conditions under which the host population gets extinct, assuming that the baseline birth rate *b* equals to one (condition leading to extinction: *μ* + *ν P* *(*λ*_0_, *β, μ, ν*) > *b*; see Appendix C).

#### Rate of neoplasia (*λ*_0_)

A high rate of neoplasia associates with a high proportion of neocancers 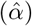 at equilibrium (Fig. 3 and 4A) and with strong selection caused by transmissible cancers 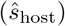 (via high cancer prevalence; Fig. 3 and 4B). While strong selection might favour sex, the high proportion of neocancers counteracts this effect, and our population genetic model predicts that this epidemiological setting as a whole does not favour the evolution of sex (Fig. 1).

On the other hand, a low rate of neoplasia associates with a low proportion of neocancers at equilibrium, which on its own would favour sex, but this also associates with transmissible cancers only causing weak selection. Here, weak selection drives the result that sex is, once again, not favoured (Fig. 1). Epidemiological considerations therefore constrain the conditions under which sex evolves: it is difficult to combine a low proportion of neocancers with strong selection in the host (Fig. S5). Overall, a low rate of neoplasia is more likely to lead to conditions favouring the evolution of sex than a high rate of neoplasia (Fig. 4C).

#### Transmission rate of cancers (*β*)

A high transmission rate of cancers associates with a low proportion of neocancers at equilibrium (Fig. 3 and 4A) and with strong selection caused by transmissible cancers (via high cancer prevalence; Fig. 3 and 4B) at equilibrium. As a result, a high transmission rate of cancers offers favourable conditions for the evolution of sex (Fig. 3 and 4C).

#### Mortality rates (*μ* and *ν*)

High mortality rates associate with a high proportion of neocancers at equilibrium (Fig. 3 and 4A). This effect occurs in our model because infected individuals dying at a rate *μ* + *ν* are replaced by susceptible newborn individuals (flow represented in purple in Fig. 2), and high mortality rates therefore associate with low cancer prevalence. The consequent low cancer prevalence reduces the risk of becoming infected by a transmitted cancer, leading to a high proportion of neocancers at equilibrium.

For high mortality rates, the consequent low cancer prevalence weakens selection caused by transmissible cancers (Equation 7). Simultaneously, however, the relative mortality rate due to cancer (*ν/μ*) directly strengthens selection caused by transmissible cancers (Equation 7). Therefore, a high baseline mortality rate (*μ*) directly weakens selection caused by transmissible cancers, in addition to its indirect effect via cancer prevalence. The situation is different in the case of a high cancer-associated mortality rate (*ν*), which has opposite direct and indirect effects on selection caused by transmissible cancers. Indeed, the high relative mortality rate due to cancer (*ν/μ*) directly strengthens selection, while the consequent low cancer prevalence weakens selection. Analytical derivations (Appendix C) show that for a high baseline mortality rate (*μ > β* − *λ*_0_), a high cancer-induced mortality rate (high *ν*) leads to strong selection. For a low baseline mortality rate (*μ* ≤ *β* − *λ*_0_), selection caused by transmissible cancers is instead maximized at intermediate values of cancer-induced mortality (intermediate *ν*) (Fig. 4B).

As a result, the combination of a low baseline mortality rate *μ* and an intermediate cancer-associated mortality rate *ν* leads to conditions favouring the evolution of sex (Fig. 4C).

As a whole, our results suggest that the conditions for transmissible cancers to select for sex are most favourable if transmissible cancers exist in a situation with a low rate of neoplasia (low *λ*_0_), a high transmission rate (high *β*) and an intermediate cancer-associated mortality rate (intermediate *ν*) in hosts with a low baseline mortality rate (low *μ*). Our epidemiological model highlights that the proportion of neocancers (*α*) and the selection coefficient caused by transmissible cancers (*s*_host_), used as independent parameters in the population genetic model, become easily linked when viewed in an epidemiological setting. Put more precisely, epidemiology constrains the conditions leading to both a low proportion of neocancers and a high selection in host (Fig. S5), since the prevalence of transmissible cancers impacts both parameters simultaneously. Our results therefore suggest that the range of parameter values providing the conditions under which sex can invade is relatively narrow (Fig. 4C). The evolution of sex is even less likely as selection on transmissible cancer due to failed infection (*s*_cancer_) decreases, regardless of other parameter values (Fig. 4C).

## DISCUSSION

At the early stages of multicellularity, when anticancer defenses were presumably less developed or prevalent than today, organisms may have been under considerable risk of transmissible cancers. At first glance, transmissible cancerous lines, akin to parasites, could induce selection on multicellular hosts to avoid infection, causing fluctuating selection and favouring the evolution of sex (Red Queen hypothesis). Nonetheless, transmissible cancers differ fundamentally from parasites: transmissible cancers are a priori genetically similar to their hosts because they originally derive from hosts’ tissues. Our formal theoretical investigation shows that antagonistic interactions between multicellular organisms and transmissible cancerous lines only rarely lead to fluctuating selection when we account for this fundamental aspect. As a result, while transmissible cancers may favour the evolution of sex in multicellular hosts under the Red Queen hypothesis, they only appear to do so under very restricted conditions.

Specifically, we investigated the implication of neoplasia – i.e., the production of cancerous lines from multicellular hosts’ tissues – for the evolution of sex under the Red Queen hypothesis. If one genotype is common in the host population, neoplasia mostly produces transmissible cancerous lines that can successfully infect this common genotype. Nonetheless, we find a striking effect of neoplasia on the coevolutionary dynamic. Neoplasia tightens the coupling between genotypic frequencies in multicellular hosts and transmissible cancers, which reduces the lag between hosts and transmissible cancers evolutionary dynamics, and thereby inhibits coevolutionary fluctuations. As the coevolutionary system reaches a stable polymorphism, fluctuating selection vanishes and sex becomes deleterious.

Our reasoning highlights that transmissible cancers share features with other processes that have been shown to inhibit the evolution of sex by dampening coevolutionary fluctuations. These include overlapping generations (Kouyos et al., 2007; but see Ashby and Gupta, 2014) and epidemiological dynamics (i.e., changes in parasite prevalence; May and Anderson, 1983; Beck, 1984; MacPherson and Otto, 2018), explaining the absence of coevolutionary cycling when we considered two types of transmissible cancers in our epidemiological model (Appendix D). Notably, parasite transmission occurring mostly among genetically similar hosts can also diminish coevolutionary cycling. It can do so by allowing the parasite population to respond more quickly to frequency changes in the hosts (Greenspoon and M’Gonigle, 2013; Greenspoon and Mideo, 2017), an effect similar to what we found for neoplasia.

In our population genetic model, the strength of selection associated with the interaction between hosts and transmissible cancers proves to be an additional important determinant of the long-term coevolutionary dynamics (see Gandon and Otto, 2007, for similar results in a standard Red Queen model). Despite the dampening effect of neoplasia, coevolutionary fluctuations may persist when host-cancer interactions associate with strong enough selection. Our population genetic model treats the proportion of transmissible cancers that recently derived from neoplasia and the selection coefficient in the host as independent parameters (following an ‘open’ approach; Cooper et al., 2018). Our complementary model with an explicit epidemiological context (a ‘closed’ approach; Cooper et al., 2018), however, shows that the values of the parameters are likely to covary in a predictable manner. Neoplasia increases the prevalence of transmissible cancers, which strengthens selection imposed by transmissible cancers. Therefore, while a low rate of neoplasia on its own would enhance coevolutionary cycling, it tends to associate with low cancer prevalence, which has the opposite effect: it weakens selection and dampens coevolutionary cycling. Since it is difficult to find conditions where both parameters take values that enhance fluctuating selection, the epidemiology of transmissible cancers as a whole constrains the conditions under which sex can evolve under the Red Queen hypothesis.

Under our main model assumptions, sexual reproduction evolves as a defensive strategy against transmissible cancers only under very restricted conditions. A slow host life history, a low rate of neoplasia, a high transmission rate, and an intermediate virulence synergistically provide the best-case scenario for fluctuating coevolutionary dynamics favouring the evolution of sex. So far, horizontally transmitted cancers have been identified in very few species in the wild, namely in dog (canine transmissible venereal tumour), in Tasmanian devil (devil facial tumour diseases), and in four bivalve species (clam leukaemia) (Ujvari et al., 2016b). In each case, phylogenetic analyses revealed that few monophyletic lines of transmissible cancerous cells widely spread in populations (Murgia et al., 2006; Pye et al., 2016; Baez-Ortega et al., 2019; Yonemitsu et al., 2019), suggesting low rates of neoplasia and high transmission rates. In two of the three cases, infection by contemporary transmissible cancers is highly virulent (in bivalve and Tasmanian devil; Barber, 2004; Lachish et al., 2007). According to our model predictions, the epidemiology of known contemporary transmissible cancers therefore matches (at least qualitatively) the restricted conditions prone to the evolution and maintenance of sex. Unfortunately, inferring the epidemiology of early transmissible cancers from those contemporary cases is speculative at best, because the nature of early transmissible cancers may have been very different when anticancer defenses were only beginning to evolve (as argued by Ujvari et al., 2016, and Thomas et al., 2019). Notably, sexual reproduction is ancestral in dogs, devils and bivalves, and as an evolutionary innovation it also precedes multicellularity (Speijer et al., 2015; Radzvilavicius and Blackstone, 2015; Weedall and Hall, 2015).

Since the focus of our model was to analyze how neoplasia affects antagonistic coevolutionary dynamics, this may come at a cost of ignoring other key features of transmissible cancers. In the following paragraphs, we therefore discuss how relaxing our modelling assumptions may change the predictions.

The life history and physiology of hosts appear to determine the epidemiology of contemporary transmissible cancers (a result somewhat akin to general ideas about cancer in short- and long-lived organisms, Kokko and Hochberg, 2015). While cancer cells are able to live freely outside of their host in bivalves (Mateo et al., 2016), the transmission of cancer cells requires physical contact between hosts in mammals (in Tasmanian devil and in dog; Murchison, 2008). In particular, transmission mainly occurs via social contacts (fights and biting), or in the case of dogs (where the tumours occur in genitalia), during copulation. Assuming that transmission requires close physical proximity among hosts, and assuming that sex cannot occur without it (a statement that excludes broadcast spawning for instance) while asexual reproduction can, then the transmission rate among sexual individuals would be particularly high (as suggested by Lehtonen et al., 2012). Similarly, while skin is generally an effective barrier against immunogenic agents, sexual behaviour may expose less protected parts of the body, enabling transmissions of cancer cells (as in the case of canine veneral tumours; Murchison, 2008). Our model did not implement any direct differences in infection dynamics between sexuals and asexuals brought about by the physics of mating. Our model effectively considered a best case scenario for the evolution of sex under the Red Queen hypothesis, where the only cost associated with sexual reproduction is the recombination load – i.e., the risk of breaking apart beneficial combinations of alleles. Although not explicitly modelled, additional costs would presumably constrain even further the conditions under which transmissible cancers can promote the evolution of sex.

Contemporary transmissible cancers have emerged and spread in a manner that allows us to track the sequence of evolutionary changes. In dog and Tasmanian devil, transmissible cancers appear to have derived from populations with low genetic diversity (Siddle et al., 2007; Miller et al., 2011; Murchison et al., 2014). More precisely, in Tasmanian devil, a transmissible cancer (Devil Facial Tumour 2) has been shown to express the host’s major histocompatibility complex class 1 (MHC1) molecules, and to match alleles that are either most widespread or non-polymorphic in the host population (Caldwell et al., 2018). Simultaneously, however, an independent but older transmissible cancer (Devil Facial Tumour 1) avoids the immune response by downregulating the expression of the MHC1 molecules (Caldwell and Siddle, 2017). Therefore, in mammals, low genetic diversity might allow newly derived transmissible cancers to persist longer in host populations, giving them time to acquire mutations necessary to infect genetically dissimilar hosts (e.g., by downregulating their immune response). This is not a process taken into account by our model. Then again, invertebrates somewhat differ from this picture as they do not possess a vertebrate-like adaptive immune system or a major histocompatibility complex, which vertebrates utilize for discriminating self from nonself. Consequently, in bivalves, transmissible cancers seem to easily infect genetically dissimilar hosts, as illustrated by cases of cross-species transmissions (Metzger et al., 2016; Yonemitsu et al., 2019).

For simplicity, our model ignores any details of how hosts reject cancers as non-self apart from a simple matching-alleles interaction among host and transmissible cancer. Real-life examples of Red Queen dynamics can be genetically rather complex: in a system of *Bacillus thuringiensis* infecting nematodes *Caenorhabditis elegans*, coevolution appears to involve copy number variations (which can evolve very rapidly) in the pathogen but many loci with small effect in the host (Papkou et al., 2019). As our model was not tailored to any particular system, our interaction model simplifies away any system-specific detail, and uses version of a self/non-self recognition system that is known to favour coevolutionary cycling in models examining the Red Queen hypothesis outside the realm of cancer (Bell and Maynard Smith, 1987; Seger and Hamilton, 1988; Howard and Lively, 1994; Agrawal and Lively, 2001, 2002). To examine more diverse multilocus settings appears a worthwhile avenue for further work. Choosing the most appropriate genetic architecture is, as a whole, a challenge because we do not know much about the allorecognition systems of the earliest multicellular organisms. It appears worthwhile to remember that if these mechanisms were incomplete (or absent), then organisms were susceptible to transmissible cancers regardless of their genetic matching with infectious cancerous cells. This scenario would appear not to promote coevolutionary fluctuations, and the evolution of sexual reproduction would, once again, not have involved the Red Queen hypothesis between hosts and cancers. However, whether this verbal argument holds for incipient (and therefore imprecise) self/nonself detection mechanisms is presently unclear. In any case, the transition to multicellularity appears to have involved regulatory changes that still play a role in cancer (Trigos et al., 2019), but whether there is also a link to sex is uncertain.

In line with the literature on the Red Queen hypothesis, we focused on the implication of fluctuating selection dynamics for the evolution of sex. Nonetheless, any antagonistic interactions can also cause ‘arm race’ dynamics characterized by the continual accumulation of adaptive mutations (Gandon et al., 2008). In such context of arm race dynamics, sexual reproduction can be beneficial by speeding up adaptation (Felsenstein, 1974). In the face of infection by transmissible cancers, sex could accelerate the development of immunity to cancerous cell lineages. More generally, by focusing on the standard Red Queen hypothesis, we did not model other evolutionary processes that can favour the evolution of sexual reproduction (except similarity selection via vertical transmission; see below).

In our main analysis, we assumed that transmissible cancers could infect any host in the population (horizontal transmission, assuming random mixing of hosts and parasites). In an additional analysis, we explored the implications of vertical transmission of cancerous cells from parent to offspring for the evolution of sexual reproduction (as advocated by Thomas et al., 2019, and as suggested by the poor recognition of fetal cells during pregnancy in mice when embryos are identical to their mother; Khosrotehrani et al., 2005), and we confirmed that this mode of transmission involves a separate mechanism, similarity selection, which can promote the evolution of sex without relying on coevolutionary fluctuations (Agrawal, 2006b). Very few studies have reported instances of vertical transmission of cancerous cells (Domazet-Lošo et al., 2014). In humans, for instance, only few cases of *in utero* cancer transmissions from mother to fetus are known (Isoda et al., 2009; Greaves and Hughes, 2018). Altogether, our results show that horizontal and vertical transmission lead to different pathways to sex. Whether the latter option still leads to sex when increasing the realism of our first model remains to be seen: we did not, for example, consider that cancer-ridden individuals might be poor at reproducing, which will reduce the prevalence of vertically transmitted cancers in the population. Threats such as late-life neocancer or being infected by a horizontally transmitted cancer may also select for other traits than sex. Life histories with risks of poor late-life performance may select for early reproduction without necessarily involving a change in reproductive mode (sexual or asexual) (as shown in the Tasmanian devil; Jones et al., 2008). While general theory for condition-dependent sex exists (and cancer, by leading to poor condition, could conceivably promote such patterns) (Hadany and Otto, 2007, 2009; Mostowy and Engelstädter, 2012; Kokko, 2020), the relative likelihood of different responses remains to be investigated theoretically.

In the metazoan hydra, asexual reproduction (‘budding’) can associate with direct vertical transmission of cancer cell, where the division of ‘parental’ cancer cells contribute to the production of newly ‘budded’ multicellular offspring (Domazet-Lošo et al., 2014). Interestingly, sexual reproduction often associates with the production of unicellular zygotes, which has been suggested to prevent the vertical transmission of non-cooperative (cancerous) cells in multicellular organisms (Grosberg and Strathmann, 1998, 2007). In brief, unicellular bottlenecks might represent an efficient way of exposing non-cooperative elements to selection early in development, hence limiting their propagation in the population. Future theoretical works could fruitfully test to what extent the transition to multicellular life, and the subsequent risk of transmitting cancer cells vertically, may have promoted the evolution of unicellular bottlenecks and of sexual reproduction (e.g., with a mechanistic model as in Pichugin et al., 2017; Staps et al., 2019). This could potentially provide a more plausible mechanism than fluctuating selection for favouring the evolution or maintenance of sexual reproduction under high cancer risk.

Overall, our theoretical models suggest that antagonistic interactions between early multicellular organisms and transmissible cancers favour the evolution of sexual reproduction as predicted under the Red Queen hypothesis only under restricted conditions. While infection by transmissible cancers causes negative frequency-dependent selection in the multicellular host, neoplasia dampens fluctuating selection (and therefore cycling coevolutionary dynamics), which underpins the Red Queen hypothesis for the evolution of sex. Nonetheless, infection by transmissible cancers could have favoured sexual reproduction via other evolutionary processes. In particular, we confirm that similarity selection caused by vertical transmission of cancerous cells could favour the evolution of sexual reproduction even without fluctuating selection.

## Acknowledgments

We thank Joanna Masel, Frédéric Thomas and one anonymous reviewer for comments that have helped improve our manuscript. This research was supported by grants from the Swiss National Science Foundation (to H.K.), from the National Institutes of Health (U54 CA217376) (to E.Y.E. and H.K.) and from the University Research Priority Program (URPP) “Evolution in Action” of the University of Zurich (to E.Y.E. and H.K.).

## References

Agrawal A. F., 2006a. Evolution of sex: why do organisms shuffle their genotypes? Current Biology, 16(17):696–704. doi: 10.1016/j.cub.2006.07.063.

Agrawal A. F., 2006b. Similarity selection and the evolution of sex: Revisiting the red queen. PLoS Biology, 4(8):1364–1371. doi: 10.1371/journal.pbio.0040265.

Agrawal A. F. and Lively C. M., 2001. Parasites and the evolution of self-fertilization. Evolution, 55(5):869–879. doi: 10.1111/j.0014-3820.2001.tb00604.x.

Agrawal A. F. and Lively C. M., 2002. Infection genetics: gene-for-gene versus matching-alleles models and all points in between. Evolutionary Ecology Research, 4:79–90.

Aktipis C. A., Boddy A. M., Jansen G., Hibner U., Hochberg M. E., Maley C. C., and Wilkinson G. S., 2015. Cancer across the tree of life: cooperation and cheating in multicellularity. Philosophical Transactions of the Royal Society B: Biological Sciences, 370(1673):20140219. doi: 10.1098/rstb.2014.0219.

Ashbel R., 1945. Spontaneous transmissible tumours in the Syrian hamster. Nature, 155(3942):607–607. doi: 10.1038/155607b0.

Ashby B. and Gupta S., 2014. Parasitic castration promotes coevolutionary cycling but also imposes a cost on sex. Evolution, 68(8):2234–2244. doi: 10.1111/evo.12425.

Baez-Ortega A., Gori K., Strakova A., Allen J. L., Allum K. M., Bansse-Issa L., Bhutia T. N., Bisson J. L., Bricenñ C., Castillo Domracheva A., Corrigan A. M., Cran H. R., Crawford J. T., Davis E., de Castro K. F., B. de Nardi A., de Vos A. P., Delgadillo Keenan L., Donelan E. M., Espinoza Huerta A. R., Faramade I. A., Fazil M., Fotopoulou E., Fruean S. N., Gallardo-Arrieta F., Glebova O., Gouletsou P. G., Häfelin Manrique R. F., Henriques J. J. G. P., Horta R. S., Ignatenko N., Kane Y., King C., Koenig D., Krupa A., Kruzeniski S. J., Kwon Y.-M., Lanza-Perea M., Lazyan M., Lopez Quintana A. M., Losfelt T., Marino G., Martínez Castañeda S., Martínez-López M. F., Meyer M., Migneco E. J., Nakanwagi B., Neal K. B., Neunzig W., Ní Leathlobhair M., Nixon S. J., Ortega-Pacheco A., Pedraza-Ordoñez F., Peleteiro M. C., Polak K., Pye R. J., Reece J. F., Rojas Gutierrez J., Sadia H., Schmeling S. K., Shamanova O., Sherlock A. G., Stammnitz M., Steenland-Smit A. E., Svitich A., Tapia Martínez L. J., Thoya Ngoka I., Torres C. G., Tudor E. M., van der Wel M. G., Vilaru B. A., Vural S. A., Walkinton O., Wang J., Wehrle-Martinez A. S., Widdowson S. A. E., Stratton M. R., Alexandrov L. B., Martincorena I., and Murchison E. P., 2019. Somatic evolution and global expansion of an ancient transmissible cancer lineage. Science, 365(6452):eaau9923. doi: 10.1126/science.aau9923.

Barber B. J., 2004. Neoplastic diseases of commercially important marine bivalves. Aquatic Living Resources, 17(4):449–466. doi: 10.1051/alr:2004052.

Barton N. H., 1995. A general model for the evolution of recombination. Genetics Research, 65(2):123–145. doi: 10.1017/s0016672300033140.

Beck K., 1984. Coevolution: mathematical analysis of host-parasite interactions. Journal of Mathematical Biology, 19(1):63–77. doi: 10.1007/BF00275931.

Bell G. 1982, The masterpiece of nature: the evolution and genetics of sexuality. University of California Press, Berkeley (CA).

Bell G. and Maynard Smith J., 1987. Short-term selection for recombination among mutually antagonistic species. Nature, 328(6125):66–68. doi: 10.1038/328066a0.

Bouvard V., Baan R., Straif K., Grosse Y., Secretan B., Ghissassi F. E., Benbrahim-Tallaa L., Guha N., Freeman C., Galichet L., and Cogliano V., 2009. A review of human carcinogens - Part B: biological agents. The Lancet Oncology, 10(4):321–322. doi: 10.1016/S1470-2045(09)70096-8.

Caldwell A. and Siddle H. V., 2017. The role of MHC genes in contagious cancer: the story of Tasmanian devils. Immunogenetics, 69(8-9): 537–545. doi: 10.1007/s00251-017-0991-9.

Caldwell A., Coleby R., Tovar C., Stammnitz M. R., Kwon Y. M., Owen R. S., Tringides M., Murchison E. P., Skjødt K., Thomas G. J., Kaufman J., Elliott T., Woods G. M., and Siddle H. V. T., 2018. The newly-arisen Devil facial tumour disease 2 (DFT2) reveals a mechanism for the emergence of a contagious cancer. eLife, 7:e35314. doi: 10.7554/eLife.35314.

Cooper G. A., Levin S. R., Wild G., and West S. A., 2018. Modeling relatedness and demography in social evolution. Evolution Letters, 2(4): 260–271. doi: 10.1002/evl3.69.

Domazet-Lošo T., Klimovich A., Anokhin B., Anton-Erxleben F., Hamm M. J., Lange C., and Bosch T. C., 2014. Naturally occurring tumours in the basal metazoan Hydra. Nature Communications, 5:4222. doi: 10.1038/ncomms5222.

Engelstädter J. and Bonhoeffer S., 2009. Red Queen dynamics with non-standard fitness interactions. PLoS Computational Biology, 5(8): e1000469. doi: 10.1371/journal.pcbi.1000469.

Feldman M. W., 1972. Selection for linkage modification: I. Random mating populations. Theoretical Population Biology, 3(3):324–346. doi: 10.1016/0040-5809(72)90007-X.

Felsenstein J., 1974. The evolutionary advantage of recombination. Genetics, 78:737–756.

Gandon S. and Otto S. P., 2007. The evolution of sex and recombination in response to abiotic or coevolutionary fluctuations in epistasis. Genetics, 175(4):1835–1853. doi: 10.1534/genetics.106.066399.

Gandon S., Buckling A., Decaestecker E., and Day T., 2008. Host-parasite coevolution and patterns of adaptation across time and space. Journal of Evolutionary Biology, 21(6):1861–1866. doi: 10.1111/j.1420-9101.2008.01598.x.

Gibson A. K., Delph L. F., Vergara D., and Lively C. M., 2018. Periodic, parasite-mediated selection for and against sex. The American Naturalist, 192(5):537–551. doi: 10.1086/699829.

Greaves M. and Hughes W., 2018. Cancer cell transmission via the placenta. Evolution, Medicine and Public Health, 2018(1):106–115. doi: 10.1093/emph/eoy011.

Greenspoon P. B. and M’Gonigle L. K., 2013. The evolution of mutation rate in an antagonistic coevolutionary model with maternal transmission of parasites. Proceedings of the Royal Society B: Biological Sciences, 280(1761):20130647. doi: 10.1098/rspb.2013.0647.

Greenspoon P. B. and M’Gonigle L. K., 2014. Host-parasite interactions and the evolution of nonrandom mating. Evolution, 68(12):3570–3580. doi: 10.1111/evo.12538.

Greenspoon P. B. and Mideo N., 2017. Parasite transmission among relatives halts Red Queen dynamics. Evolution, 71(3):747–755. doi: 10.1111/evo.13157.

Grosberg R. K. and Strathmann R. R., 1998. One cell, two cell, red cell, blue cell: the persistence of a unicellular stage in multicellular life histories. Trends in Ecology and Evolution, 13(3):112–116. doi: 10.1016/S0169-5347(97)01313-X.

Grosberg R. K. and Strathmann R. R., 2007. The evolution of multicellularity: a minor major transition? Annual Review of Ecology, Evolution, and Systematics, 38(1):621–654. doi: 10.1146/annurev.ecolsys.36.102403.114735.

Hadany L. and Otto S. P., 2007. The evolution of condition-dependent sex in the face of high costs. Genetics, 176(3):1713–1727. doi: 10.1534/genetics.107.074203.

Hadany L. and Otto S. P., 2009. Condition-dependent sex and the rate of adaptation. American Naturalist, 174:S71–S78. doi: 10.1086/599086.

Haldane J. B. S., 1949. Disease and evolution. La Ricerca Scientifica, 19:68–76.

Hamilton W. D., 1975. Gamblers since life began: barnacles, aphids, elms. The Quarterly Review of Biology, 50(2):175–180. doi: 10.1086/408439.

Hamilton W. D., 1980. Sex versus non-sex versus parasite. Oikos, 35(2):282–290. doi: 10.2307/3544435.

Hartfield M. and Keightley P. D., 2012. Current hypotheses for the evolution of sex and recombination. Integrative Zoology, 7(2):192–209. doi: 10.1111/j.1749-4877.2012.00284.x.

Howard R. S. and Lively C. M., 1994. Parasitism, mutation accumulation and the maintenance of sex. Nature, 367(6463):554–557. doi: 10.1038/367554a0.

Isoda T., Ford A. M., Tomizawa D., Van Delft F. W., De Castro D. G., Mitsuiki N., Score J., Taki T., Morio T., Takagi M., Saji H., Greaves M., and Mizutani S., 2009. Immunologically silent cancer clone transmission from mother to offspring. Proceedings of the National Academy of Sciences of the United States of America, 106(42):17882–17885. doi: 10.1073/pnas.0904658106.

Jones M. E., Cockburn A., Hamede R., Hawkins C., Hesterman H., Lachish S., Mann D., McCallum H., and Pemberton D., 2008. Life-history change in disease-ravaged Tasmanian devil populations. Proceedings of the National Academy of Sciences of the United States of America, 105(29):10023–10027. doi: 10.1073/pnas.0711236105.

Khosrotehrani K., Johnson K. L., Guégan S., Stroh H., and Bianchi D. W., 2005. Natural history of fetal cell microchimerism during and following murine pregnancy. Journal of Reproductive Immunology, 66(1):1–12. doi: 10.1016/j.jri.2005.02.001.

Kokko H., 2020. When synchrony makes the best of both worlds even better: How well do we really understand facultative sex? The American Naturalist, 195(2):380–392. doi: 10.1086/706812.

Kokko H. and Hochberg M. E., 2015. Towards cancer-aware life-history modelling. Philosophical Transactions of the Royal Society B: Biological Sciences, 370(1673):20140234. doi: 10.1098/rstb.2014.0234.

Kouyos R. D., Salathé M., and Bonhoeffer S., 2007. The Red Queen and the persistence of linkage-disequilibrium oscillations in finite and infinite populations. BMC Evolutionary Biology, 7:211. doi: 10.1186/1471-2148-7-211.

Lachish S., Jones M., and McCallum H., 2007. The impact of disease on the survival and population growth rate of the Tasmanian devil. Journal of Animal Ecology, 76(5):926–936. doi: 10.1111/j.1365-2656.2007.01272.x.

Lehtonen J., Jennions M. D., and Kokko H., 2012. The many costs of sex. Trends in Ecology and Evolution, 27(3):172–178. doi: 10.1016/j.tree.2011.09.016.

Levin D. A., 1975. Pest pressure and recombination systems in plants. The American Naturalist, 109(968):437–451. doi: 10.1086/283012.

Lively C. M., 2010. A review of Red Queen models for the persistence of obligate sexual reproduction. Journal of Heredity, 101:13–20. doi: 10.1093/jhered/esq010.

Lively C. M. and Dybdahl M. F., 2000. Parasite adaptation to locally common host genotypes. Nature, 405(6787):679–681. doi: 10.1038/35015069.

MacPherson A. and Otto S. P., 2018. Joint coevolutionary-epidemiological models dampen Red Queen cycles and alter conditions for epidemics. Theoretical Population Biology, 122:137–148. doi: 10.1016/j.tpb.2017.12.003.

Mateo D. R., MacCallum G. S., and Davidson J., 2016. Field and laboratory transmission studies of haemic neoplasia in the soft-shell clam, Mya arenaria, from Atlantic Canada. Journal of Fish Diseases, 39(8):913–927. doi: 10.1111/jfd.12426.

May R. M. and Anderson R. M., 1983. Epidemiology and genetics in the coevolution of parasites and hosts. Proceedings of the Royal Society B: Biological Sciences, 219(1216):281–313. doi: 10.1098/rspb.1983.0075.

Metzger M. J., Reinisch C., Sherry J., and Goff S. P., 2015. Horizontal transmission of clonal cancer cells causes leukemia in soft-shell clams. Cell, 161(2):255–263. doi: 10.1016/j.cell.2015.02.042.

Metzger M. J., Villalba A., Carballal M. J., Iglesias D., Sherry J., Reinisch C., Muttray A. F., Baldwin S. A., and Goff S. P., 2016. Widespread transmission of independent cancer lineages within multiple bivalve species. Nature, 534(7609):705–709. doi: 10.1038/nature18599.

Miller W., Hayes V. M., Ratan A., Petersen D. C., Wittekindt N. E., Miller J., Walenz B., Knight J., Qi J., Zhao F., Wang Q., Bedoya-Reina O. C., Katiyar N., Tomsho L. P., Kasson L. M., Hardie R.-A., Woodbridge P., Tindall E. A., Bertelsen M. F., Dixon D., Pyecroft S., Helgen K. M., Lesk A. M., Pringle T. H., Patterson N., Zhang Y., Kreiss A., Woods G. M., Jones M. E., and Schuster S. C., 2011. Genetic diversity and population structure of the endangered marsupial Sarcophilus harrisii (Tasmanian devil). Proceedings of the National Academy of Sciences of the United States of America, 108(30):12348–12353. doi: 10.1073/pnas.1102838108.

Miroshnychenko D., Baratchart E., Ferrall-Fairbanks M. C., Velde R. V., Laurie M. A., Bui M. M., Altrock P. M., Basanta D., and Marusyk A., 2020. Spontaneous cell fusions as a mechanism of parasexual recombination in tumor cell populations. BioRxiv preprint. doi: 10.1101/2020.03.09.984419.

Mostowy R. and Engelstädter J., 2012. Host-parasite coevolution induces selection for condition-dependent sex. Journal of Evolutionary Biology, 25(10):2033–2046. doi: 10.1111/j.1420-9101.2012.02584.x.

Murchison E. P., 2008. Clonally transmissible cancers in dogs and Tasmanian devils. Oncogene, 27(S2):S19–S30. doi: 10.1038/onc.2009.350.

Murchison E. P., Wedge D. C., Alexandrov L. B., Fu B., Martincorena I., Ning Z., Tubio J. M. C., Werner E. I., Allen J., De Nardi A. B., Donelan E. M., Marino G., Fassati A., Campbell P. J., Yang F., Burt A., Weiss R. A., and Stratton M. R., 2014. Transmissible dog cancer genome reveals the origin and history of an ancient cell Lineage. Science, 343(6169):437–440. doi: 10.1126/science.1247167.

Murgia C., Pritchard J. K., Kim S. Y., Fassati A., and Weiss R. A., 2006. Clonal origin and evolution of a transmissible cancer. Cell, 126(3): 477–487. doi: 10.1016/j.cell.2006.05.051.

Nee S., 1989. Antagonistic co-evolution and the evolution of genotypic randomization. Journal of Theoretical Biology, 140(4):499–518. doi: 10.1016/S0022-5193(89)80111-0.

Otto S. P. and Lenormand T., 2002. Resolving the paradox of sex and recombination. Nature Reviews Genetics, 3(4):252–261. doi: 10.1038/nrg761.

Otto S. P. and Michalakis Y., 1998. The evolution of recombination in changing environments. Trends in Ecology & Evolution, 13(4):145–151. doi: 10.1016/S0169-5347(97)01260-3.

Otto S. P. and Nuismer S. L., 2004. Species interactions and the evolution of sex. Science, 304(5673):1018–1020. doi: 10.1126/science.1094072.

Papkou A., Guzella T., Yang W., Koepper S., Pees B., Schalkowski R., Barg M.-C., Rosenstiel P. C., Teotónio H., and Schulenburg H., 2019. The genomic basis of Red Queen dynamics during rapid reciprocal hostpathogen coevolution. Proceedings of the National Academy of Sciences of the United States of America, 116(3):923–928. doi: 10.1073/pnas.1810402116.

Pichugin Y., Peña J., Rainey P. B., and Traulsen A., 2017. Fragmentation modes and the evolution of life cycles. PLoS Computational Biology, 13(11):e1005860. doi: 10.1371/journal.pcbi.1005860.

Pienta K. J., Hammarlund E. U., Axelrod R., Brown J. S., and Amend S. R., 2020. Poly-aneuploid cancer cells promote evolvability, generating lethal cancer. Evolutionary Applications, in press. doi: 10.1111/eva.12929.

Pye R. J., Pemberton D., Tovar C., Tubio J. M. C., Dun K. A., Fox S., Darby J., Hayes D., Knowles G. W., Kreiss A., Siddle H. V. T., Swift K., Lyons A. B., Murchison E. P., and Woods G. M., 2016. A second transmissible cancer in Tasmanian devils. Proceedings of the National Academy of Sciences, 113(2):374–379. doi: 10.1073/pnas.1519691113.

Radzvilavicius A. L. and Blackstone N. W., 2015. Conflict and cooperation in eukaryogenesis: implications for the timing of endosymbiosis and the evolution of sex. Journal of the Royal Society Interface, 12(111):20150584. doi: 10.1098/rsif.2015.0584.

Salathé M., Kouyos R. D., Regoes R. R., and Bonhoeffer S., 2008. Rapid parasite adaptation drives selection for high recombination rates. Evolution, 62(2):295–300. doi: 10.1111/j.1558-5646.2007.00265.x.

Salathé M., Kouyos R. D., and Bonhoeffer S., 2009. On the causes of selection for recombination underlying the Red Queen hypothesis. The American Naturalist, 174(S1):S31–S42. doi: 10.1086/599085.

Schwander T., Marais G., and Roze D., 2014. Sex uncovered: the evolutionary biology of reproductive systems. Journal of Evolutionary Biology, 27(7):1287–1291. doi: 10.1111/jeb.12424.

Seger J. and Hamilton W. D. Parasites and sex. In Michod R. E. and Levin B. R., editors, The Evolution of Sex: An Examination of Current Ideas, pages 176–193. Sunderland, sinauer as edition, 1988.

Siddle H. V., Kreiss A., Eldridge M. D. B., Noonan E., Clarke C. J., Pyecroft S., Woods G. M., and Belov K., 2007. Transmission of a fatal clonal tumor by biting occurs due to depleted MHC diversity in a threatened carnivorous marsupial. Proceedings of the National Academy of Sciences, 104(41):16221–16226. doi: 10.1073/pnas.0704580104.

Sidow A. and Spies N., 2015. Concepts in solid tumor evolution. Trends in Genetics, 31(4):208–214. doi: 10.1016/j.tig.2015.02.001.

Speijer D., Lukeš J., and Eliáš M., 2015. Sex is a ubiquitous, ancient, and inherent attribute of eukaryotic life. Proceedings of the National Academy of Sciences of the United States of America, 112(29):8827–8834. doi: 10.1073/pnas.1501725112.

Staps M., van Gestel J., and Tarnita C. E., 2019. Emergence of diverse life cycles and life histories at the origin of multicellularity. Nature Ecology and Evolution, 3(8):1197–1205. doi: 10.1038/s41559-019-0940-0.

Thomas F., Madsen T., Giraudeau M., Misse D., Hamede R., Vincze O., Renaud F., Roche B., and Ujvari B., 2019. Transmissible cancer and the evolution of sex. PLoS Biology, 17(6):e3000275. doi: 10.1371/journal.pbio.3000275.

Trigos A. S., Pearson R. B., Papenfuss A. T., and Goode D. L., 2019. Somatic mutations in early metazoan genes disrupt regulatory links between unicellular and multicellular genes in cancer. eLife, 8:e40947. doi: 10.7554/eLife.40947.

Ujvari B., Gatenby R. A., and Thomas F., 2016a. Transmissible cancers, are they more common than thought? Evolutionary Applications, 9(5):633–634. doi: 10.1111/eva.12372.

Ujvari B., Gatenby R. A., and Thomas F., 2016b. The evolutionary ecology of transmissible cancers. Infection, Genetics and Evolution, 39: 293–303. doi: 10.1016/j.meegid.2016.02.005.

Van Valen L., 1973. A new evolutionary law. Evolutionary Theory, 1:1–30.

Weedall G. D. and Hall N., 2015. Sexual reproduction and genetic exchange in parasitic protists. Parasitology, 142(S1):S120–S127. doi: 10.1017/S0031182014001693.

Yonemitsu M. A., Giersch R. M., Polo-Prieto M., Hammel M., Simon A., Cremonte F., Avilés F. T., Merino-Véliz N., Burioli E. A., Muttray A. F., Sherry J., Reinisch C., Baldwin S. A., Goff S. P., Houssin M., Arriagada G., Vazquez N., Bierne N., and Metzger M. J., 2019. A single clonal lineage of transmissible cancer identified in two marine mussel species in South America and Europe. eLife, 8:e47788. doi: 10.7554/eLife.47788.

