## Supplementary Information for "Transmissible cancers and the evolution of sex under the Red Queen hypothesis"

##### Contents

Pages 2 - 5: Supplementary Figures S1-S5

Pages 6 - 9: Appendix A: Analytical derivation of a simplified one-locus population genetic model

Pages 10 - 11: Appendix B: Numerical analysis of a population genetic model with similarity selection

Pages 12 - 21: Appendix C: Analytical derivation of the epidemiological model

Pages 22 - 24: Appendix D: Numerical analysis of an epidemiological model with two types of cancers

Page 25: References

### SUPPLEMENTARY FIGURES

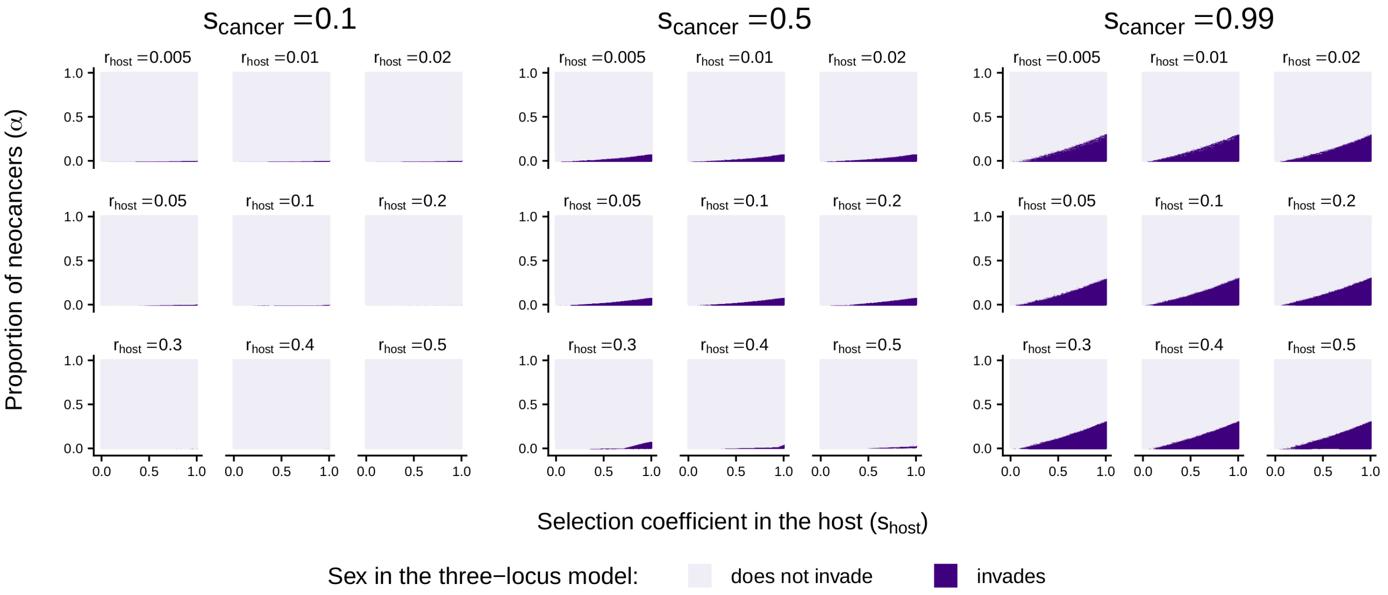


Figure S1: Evolution of sex associated to different recombination rates ($r_{\text{host}}$). Sensitivity of the three-locus population genetic model to the selection coefficients ($s_{\text{host}}, s_{\text{cancer}}$) and to the proportion of transmissible neocancers deriving from hosts' neoplasms ($\alpha$). The conditions under which sex can invade are more restricted when sex associates with a high recombination rate (leading to a high recombination load). Sex without genetic mixing is neutral compared to asexual reproduction (i.e., if $r_{\text{host}} = 0$ in our haploid case, not shown). Therefore, sex is more strongly favoured if it associates with an intermediate recombination rate $r_{\text{host}}$. See Figure 1 for more details.

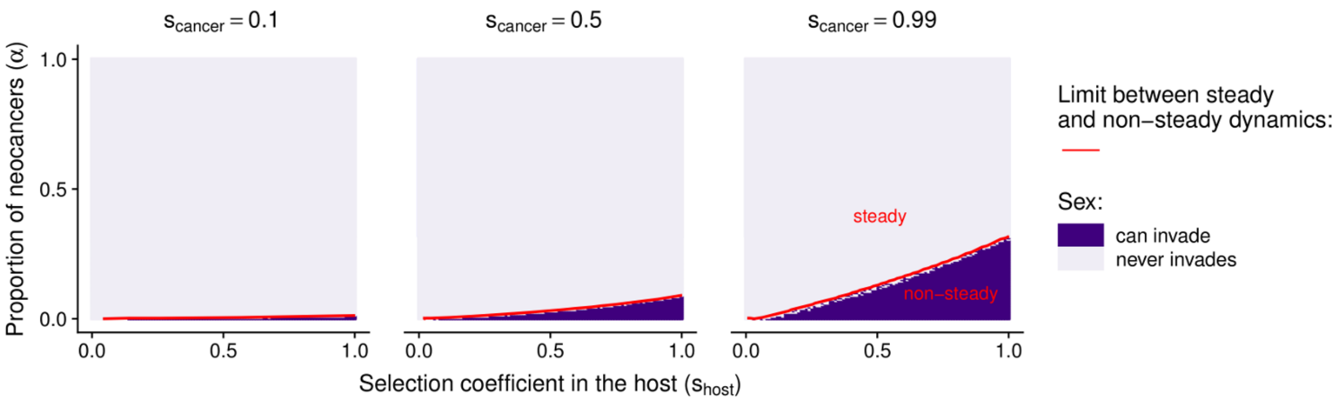


Figure S2: Coevolutionary dynamics between hosts and transmissible cancers, and evolution from facultative sex (50% asexual reproduction, and 50% sexual reproduction) to obligate sex (100% sexual reproduction). Sensitivity of the population genetic models to the selection coefficients ($s_{\mathrm{host}}, s_{\mathrm{cancer}}$) and to the proportion of transmissible neocancers that are recently derived from the original host ($\alpha$). Red lines delimit the parameter spaces leading to non-steady and steady coevolutionary dynamics. The dynamic is defined as 'steady' when the variance in genotypic frequencies over 500 time steps is below $10^{-10}$. Dark purple indicates conditions under which a modifier allele associated with obligate sexual reproduction (and with recombination, at least for one of the recombination rates tested) can invade in at least one of the 100 simulation runs. We get the same results as in Figure 1.

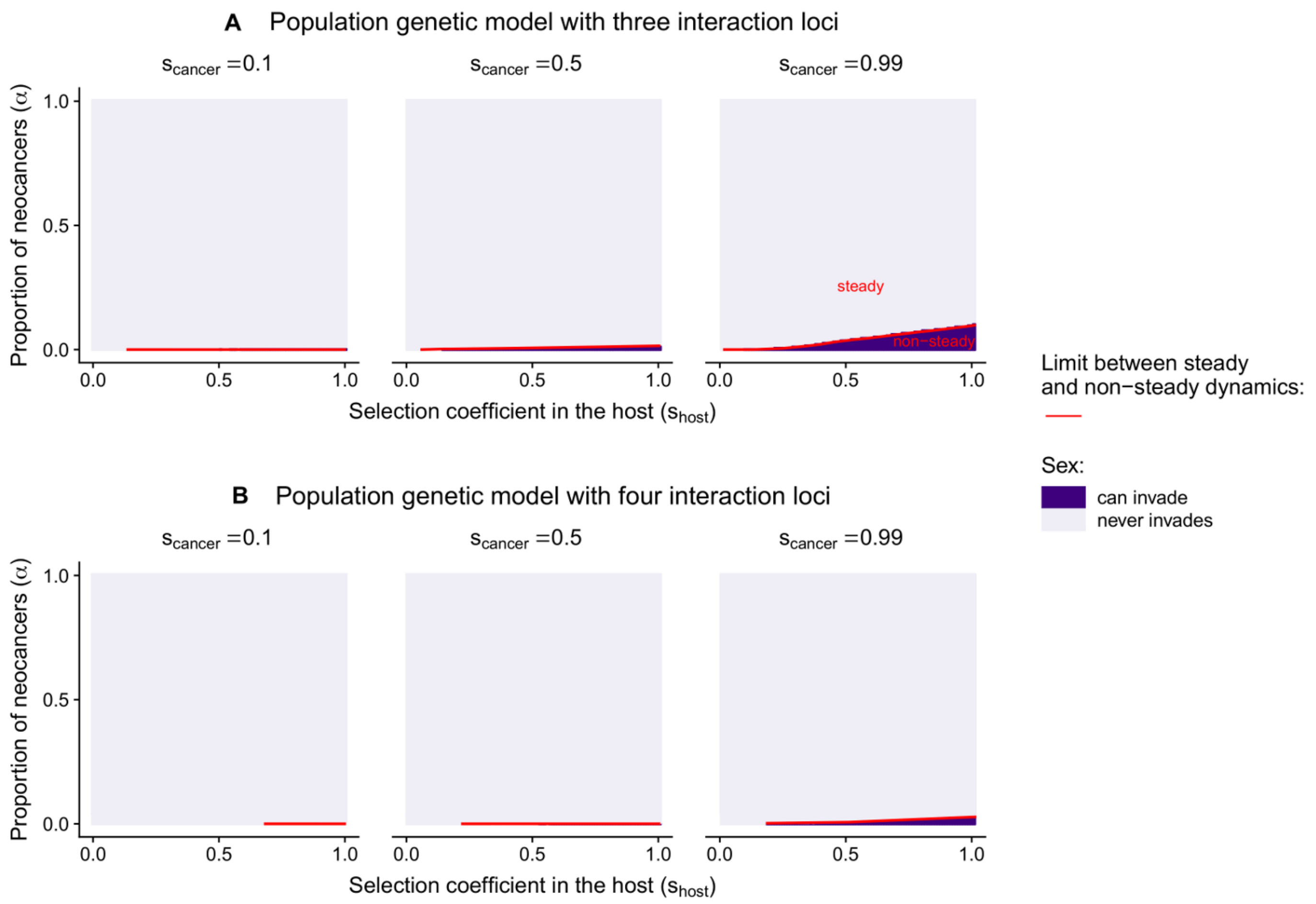


Figure S3: Coevolutionary dynamics between hosts and transmissible cancers, and evolution of sex in population genetic models with three or four interaction loci (determining the outcome of the interaction between hosts and cancers). Sensitivity of the population genetic models to the selection coefficients ($s_{\text{host}}, s_{\text{cancer}}$) and to the proportion of transmissible neocancers that are recently derived from the original host ($\alpha$). Red lines delimit the parameter spaces leading to non-steady and steady coevolutionary dynamics. The dynamic is defined as 'steady' when the variance in genotypic frequencies over 500 time steps is below $10^{-10}$. Dark purple indicates conditions under which a modifier allele associated with sexual reproduction (and with recombination, at least for one of the recombination rates tested) can invade in at least one of the 100 simulation runs. Sex is favoured mostly within a restricted genotypic space under the Red Queen hypothesis (as shown in Otto and Nuismer, 2004) (but see Iles et al. (2003) and Hickey and Golding (2018) accounting for other evolutionary processes favouring sexual reproduction). Additionally, neoplasia dampens coevolutionary cycling even when considering more than two interaction loci.

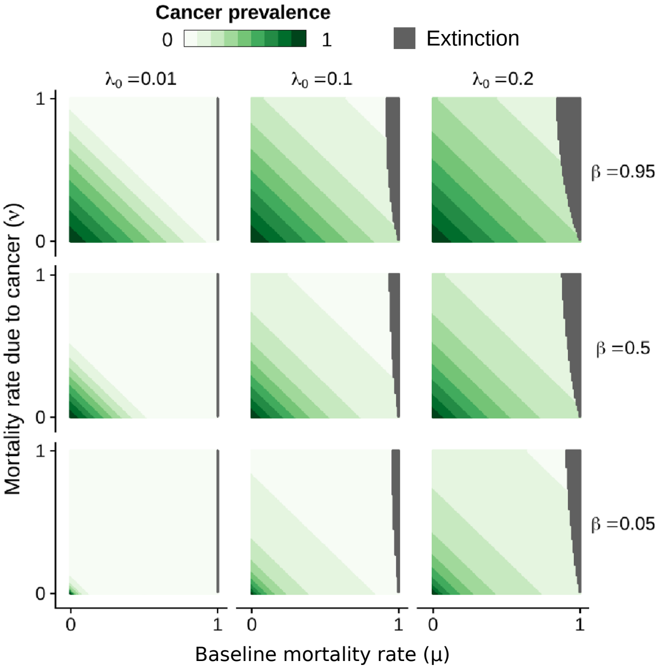


Figure S4: Prevalence $P^*$ of transmissible cancers at equilibrium. A high prevalence associates with a high selection coefficient in the host (high $\hat{s}_{\text{host}}$, Fig. 4B). In gray, we represent the conditions under which the host population gets extinct, assuming that the baseline birth rate $b$ equals to one (condition leading to extinction: $\mu + \nu\, P^*(\lambda_0, \beta, \mu, \nu) > b$; see Appendix C).

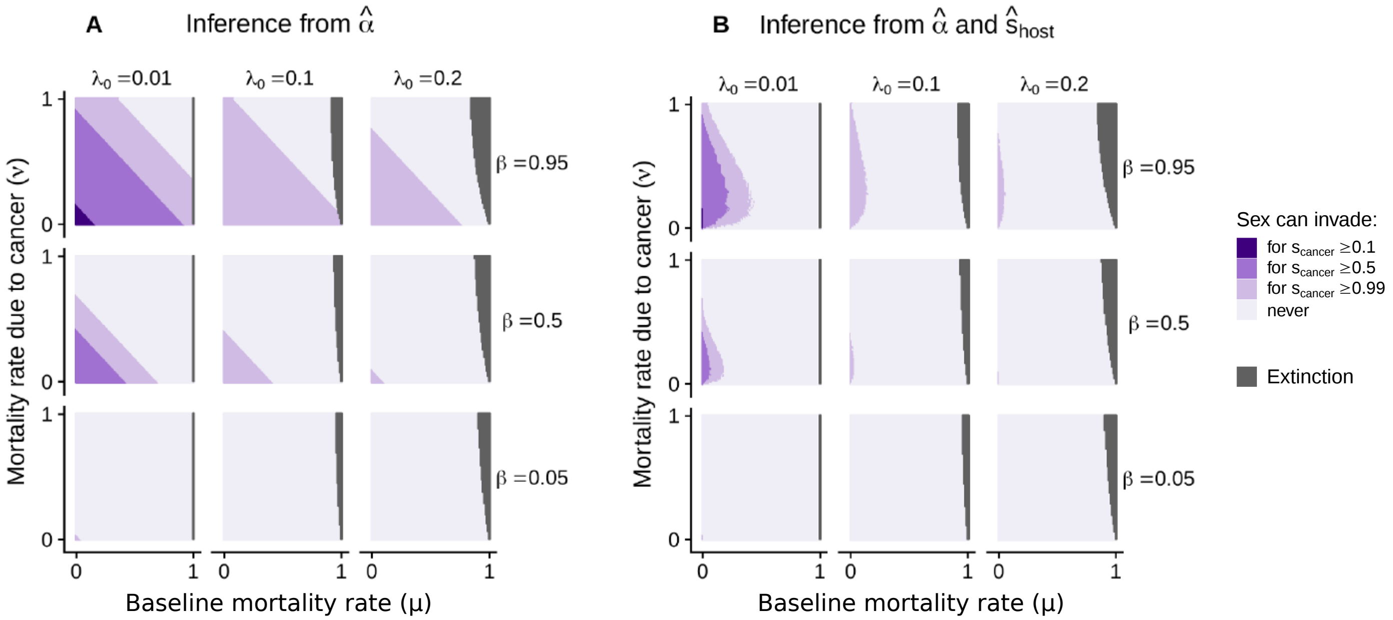


Figure S5: Conditions under which sex can invade (inferred from the three-locus population genetic model) in the epidemiological model. At equilibrium, we determine the values of $(\hat{\alpha}, \hat{s}_{\text{host}})$, and the conditions favouring the evolution of sex are inferred from $\hat{\alpha}$ only (A) or from $(\hat{\alpha}, \hat{s}_{\text{host}})$ (B). Decreased rate of neoplasia ($\lambda_0$) leads to the conditions of $\hat{\alpha}$ that are prone to the evolution of sex (A). Nonetheless, it also associates with a decrease in the selection coefficient caused by transmissible cancers (Fig. 4B), thereby inhibiting the evolution of sex (B). In gray, we represent the conditions under which the host population gets extinct, assuming that the baseline birth rate $b$ equals to one (condition leading to extinction: $\mu + \nu\, P^*(\lambda_0, \beta, \mu, \nu) > b$; see Appendix C). See Figure 1 for more details.

### APPENDIX A: Analytical derivation of a simplified one-locus population genetic model

#### Purpose

To gain further insights into the phenomenon presented in the main manuscript, we consider a simplified one-locus population genetic model and solve it analytically.

Hosts and transmissible cancers are haploid. This simplified population genetic model is based on a single autosomal haploid locus $A$ with two possible alleles (A/a), controlling the interaction between hosts and transmissible cancers (with selection coefficients $s_{\text{host}}$ in case of a match, and $s_{\text{cancer}}$ in case of a mismatch, respectively). Time is discrete, and we assume that a proportion of transmissible cancers $\alpha$ derives from neocancers.

Contrary to the population genetic model analyzed numerically in the manuscript, this model does not include a modifier locus controlling the reproduction mode of the host. Here, our aim is to investigate analytically the conditions under which coevolutionary cycling (so-called 'Red Queen dynamics') is taking place between hosts and transmissible cancers. Likewise, we assume there is no mutation.

Below, we show that all equilibrium points are unstable for:

$$\boxed{\alpha < \frac{s_{\text{host}}\, s_{\text{cancer}}}{(2 - s_{\text{host}})(2 - s_{\text{cancer}}) + s_{\text{host}}\, s_{\text{cancer}}}} \tag{1}$$

Under this condition, the long-term behaviour of the system is not steady. We cannot show analytically that this corresponds to a limit cycle; it could also be chaotic dynamics. In both cases, such non-steady long-term behaviour gives scope for the evolution of sex under the Red Queen Hypothesis.

#### Recursive equations

The systems of equations describing the changes in frequency of allele A in the host and the transmissible cancers ($f_{\text{h}}$ and $f_{\text{c}}$) over one generation are:

$$\begin{cases} f'_{\text{h}} &= \dfrac{f_{\text{h}}\,\left[\,(1 - s_{\text{host}})\; f^*_{\text{c}} + (1 - f^*_{\text{c}})\,\right]}{f_{\text{h}}\,\left[\,(1 - s_{\text{host}})\; f^*_{\text{c}} + (1 - f^*_{\text{c}})\,\right] + (1 - f_{\text{h}})\,\left[\,f^*_{\text{c}} + (1 - s_{\text{host}})\;(1 - f^*_{\text{c}})\,\right]} \\ f'_{\text{c}} &= \dfrac{f^*_{\text{c}}\,\left[\,f_{\text{h}} + (1 - s_{\text{cancer}})\;(1 - f_{\text{h}})\,\right]}{f^*_{\text{c}}\,\left[\,f_{\text{h}} + (1 - s_{\text{cancer}})\;(1 - f_{\text{h}})\,\right] + (1 - f^*_{\text{c}})\,\left[\,(1 - s_{\text{cancer}})\; f_{\text{h}} + (1 - f_{\text{h}})\,\right]} \end{cases} \tag{2}$$

With

$$f^*_{\text{c}} = \alpha\, f_{\text{h}} + (1 - \alpha)\; f_{\text{c}} \tag{3}$$

Parameter $\alpha$ corresponds to the proportion of transmissible cancers that derive from neocancers. Hosts suffer a fitness cost if they get infected by cancers with the same genotype than their own (fitness coefficient $1 - s_{\text{host}}$). Transmissible cancers suffer a fitness cost if they infect hosts with a different genotype than their own (fitness coefficient $1 - s_{\text{cancer}}$).

This system of equations can be simplified as:

$$\begin{cases} f'_{\text{h}} &= \dfrac{f_{\text{h}}\,\left[\,1 - s_{\text{host}}\, f^*_{\text{c}}\,\right]}{1 + s_{\text{host}}\,[f_{\text{h}} + f^*_{\text{c}} - 1 - 2\, f_{\text{h}}\, f^*_{\text{c}}]} \\ f'_{\text{c}} &= \dfrac{f^*_{\text{c}}\,\left[\,1 - s_{\text{cancer}}\;(1 - f_{\text{h}})\,\right]}{1 + s_{\text{cancer}}\,[2\, f_{\text{h}}\, f^*_{\text{c}} - f_{\text{h}} - f^*_{\text{c}}]} \end{cases} \tag{4}$$

With $0 \leq \alpha \leq 1$, $0 < s_{\text{host}} < 1$ and $0 < s_{\text{cancer}} < 1$.

### Equilibria

This system of equations is characterized by three or five equilibrium points depending on parameter $\alpha$:

- Fixation of different alleles in host and cancer: $(f_{\mathrm{h}}, f_{\mathrm{c}}) = (1,0)$ or $(0,1)$ **only if** $\boldsymbol{\alpha = 0}$
- Fixation of the same allele in host and cancer: $(f_{\mathrm{h}}, f_{\mathrm{c}}) = (0,0)$ or $(1,1)$
- Equal frequencies of the two alleles in both host and cancer: $(f_{\mathrm{h}}, f_{\mathrm{c}}) = (0.5, 0.5)$

### Local stability

At each equilibrium point, the local stability can be inferred from the eigenvalues of the Jacobian matrix $\mathbf{J}$. The complete expression of $\mathbf{J}$ is very unwieldy. Below, we only show the expressions of the Jacobian matrices evaluated at the equilibrium points.

#### 1. Fixation of different alleles (if $\alpha = 0$)

Around the equilibrium points $(0,1)$ and $(1,0)$, the expression of the Jacobian matrix is:

$$\mathbf{J}_{(0,1),(1,0)|\alpha=0} = \begin{bmatrix} 1 - s_{\mathrm{host}} & 0 \\ 0 & \frac{1}{1-s_{\mathrm{cancer}}} \end{bmatrix} \tag{5}$$

Given that the eigenvalue $\frac{1}{1-s_{\mathrm{cancer}}} > 1$ (on the diagonal), those equilibrium points are always locally **unstable**.

#### 2. Fixation of the same allele

Around the equilibrium points $(0,0)$ and $(1,1)$, the expression of the Jacobian matrix is:

$$\mathbf{J}_{(0,0),(1,1)} = \begin{bmatrix} \frac{1}{1-s_{\mathrm{host}}} & 0 \\ \alpha\,(1 - s_{\mathrm{cancer}}) & (1 - s_{\mathrm{cancer}})(1-\alpha) \end{bmatrix} \tag{6}$$

Given that the eigenvalue $\frac{1}{1-s_{\mathrm{host}}} > 1$ (on the diagonal), those equilibrium points are always locally **unstable**.

#### 3. Equal frequencies of the two alleles

Around the equilibrium points $(0.5, 0.5)$, the expression of the Jacobian matrix is:

$$\mathbf{J}^* = \begin{bmatrix} 1 - \frac{\alpha\, s_{\mathrm{host}}}{2-s_{\mathrm{host}}} & \frac{-s_{\mathrm{host}}\,(1-\alpha)}{2-s_{\mathrm{host}}} \\ \alpha + \frac{s_{\mathrm{cancer}}}{2-s_{\mathrm{cancer}}} & 1-\alpha \end{bmatrix} \tag{7}$$

We can show that this equilibrium point is unstable for:

$$\alpha < \frac{s_{\mathrm{host}}\, s_{\mathrm{cancer}}}{(2 - s_{\mathrm{host}})(2 - s_{\mathrm{cancer}}) + s_{\mathrm{host}}\, s_{\mathrm{cancer}}} \tag{8}$$

***Proof:***

The trace and determinant of $\mathbf{J}^*$ are:

$$\mathrm{trace}\,(\mathbf{J}^*) = 2 - \alpha - \frac{\alpha\, s_{\mathrm{host}}}{2 - s_{\mathrm{host}}} \tag{9}$$

$$\det\,(\mathbf{J}^*) = \frac{(1-\alpha)\ \left[(2 - s_{\mathrm{cancer}})\,(2 - s_{\mathrm{host}}\,(1+\alpha)) + s_{\mathrm{host}}\,(\alpha\,(2 - s_{\mathrm{cancer}}) + s_{\mathrm{cancer}})\right]}{(2 - s_{\mathrm{host}})(2 - s_{\mathrm{cancer}})} \tag{10}$$

The eigenvalues can be expressed as:

$$\lambda = \frac{\mathrm{trace}\,(\mathbf{J}^*) \pm \sqrt{\mathrm{trace}\,(\mathbf{J}^*)^2 - 4\det\,(\mathbf{J}^*)}}{2} \tag{11}$$

The eigenvalues are either complex or real, depending on the sign of:

$$\text{trace}\left(\mathbf{J}^*\right)^2 - 4\det\left(\mathbf{J}^*\right) = \frac{4\left[(2-s_{\text{cancer}})\,\alpha^2 - 2\,s_{\text{host}}\,(1-s_{\text{cancer}})\,(2-s_{\text{host}})\,\alpha - s_{\text{host}}\,s_{\text{cancer}}\,(2-s_{\text{host}})\right]}{(s_{\text{host}}-2)^2(2-s_{\text{cancer}})} \tag{12}$$

which has the same sign as the polynomial function:

$$P(\alpha) = (2-s_{\text{cancer}})\,\alpha^2 - 2\,s_{\text{host}}\,(1-s_{\text{cancer}})\,(2-s_{\text{host}})\,\alpha - s_{\text{host}}\,s_{\text{cancer}}\,(2-s_{\text{host}}) \tag{13}$$

The discriminant of this polynomial function is positive:

$$\Delta_P = 4\,s_{\text{host}}^2\,(1-s_{\text{cancer}})^2\,(2-s_{\text{host}})^2 + 4\,(2-s_{\text{cancer}})\,(2-s_{\text{host}})\,s_{\text{host}}\,s_{\text{cancer}} \tag{14}$$

The product of the roots of this polynomial function is negative (Vieta's formula). Therefore, this polynomial function is characterized by a single positive root $A_+$:

$$A_+ = \frac{2\,s_{\text{host}}\,(1-s_{\text{cancer}})\,(2-s_{\text{host}}) + \sqrt{\Delta_P}}{2\,(2-s_{\text{cancer}})} \tag{15}$$

Therefore, $\text{trace}\left(\mathbf{J}^*\right)^2 - 4\det\left(\mathbf{J}^*\right) > 0$ if $\alpha > A_+$. This means that the eigenvalues are real for $\alpha \geq A_+$, and complex for $\alpha < A_+$. We now infer the stability of the equilibrium point under these two conditions (points a and b below).

*a) For $\alpha \geq A_+$:* Eigenvalues are real. Stability requires that the absolute value of the leading eigenvalue $\lambda_{\text{L}}$ is less than one. Given that $\text{trace}\left(\mathbf{J}^*\right) > 0$, the leading eigenvalue is:

$$\lambda_{\text{L}} = \frac{\text{trace}\left(\mathbf{J}^*\right) + \sqrt{\text{trace}\left(\mathbf{J}^*\right)^2 - 4\det\left(\mathbf{J}^*\right)}}{2} > 0 \tag{16}$$

We can show that $\lambda_{\text{L}} < 1$ by determining the sign of

$$\lambda_{\text{L}} - 1 = \frac{\sqrt{\text{trace}\left(\mathbf{J}^*\right)^2 - 4\det\left(\mathbf{J}^*\right)} - \alpha\left(1 + \frac{s_{\text{host}}}{2-s_{\text{host}}}\right)}{2} \tag{17}$$

Which has the same sign as:

$$\text{trace}\left(\mathbf{J}^*\right)^2 - 4\det\left(\mathbf{J}^*\right) - \alpha^2\left(1 + \frac{s_{\text{host}}}{2-s_{\text{host}}}\right)^2 = \frac{4\,s_{\text{host}}\,(-2\alpha\,(1-s_{\text{cancer}}) - s_{\text{cancer}})}{(1-s_{\text{cancer}})(1-s_{\text{host}})} < 0 \tag{18}$$

Therefore, $\lambda_{\text{L}} < 1$, meaning that the equilibrium point is stable.

*b) For $\alpha < A_+$:* Eigenvalues are complex, and take the form:

$$\lambda = \frac{\text{trace}\left(\mathbf{J}^*\right)}{2} \pm i\frac{\sqrt{-\text{trace}\left(\mathbf{J}^*\right)^2 + 4\det\left(\mathbf{J}^*\right)}}{2} \tag{19}$$

Stability requires that:

$$\sqrt{\left(\frac{\text{trace}\left(\mathbf{J}^*\right)}{2}\right)^2 + \left(\frac{\sqrt{-\text{trace}\left(\mathbf{J}^*\right)^2 + 4\det\left(\mathbf{J}^*\right)}}{2}\right)^2} = \sqrt{\det\left(\mathbf{J}^*\right)} < 1 \tag{20}$$

Given that $\sqrt{\det\left(\mathbf{J}^*\right)} > 0$, this means that stability requires $\det\left(\mathbf{J}^*\right) < 1$:

$$\det\left(\mathbf{J}^*\right) - 1 = \frac{-\alpha\,(2-s_{\text{host}})(2-s_{\text{cancer}}) - \alpha\,s_{\text{host}}\,s_{\text{cancer}} + s_{\text{host}}\,s_{\text{cancer}}}{(2-s_{\text{host}})(2-s_{\text{cancer}})} \tag{21}$$

Therefore, stability requires that $\alpha$ is higher than a threshold value $A^*$:

$$\alpha > A^* = \frac{s_{\text{host}}\,s_{\text{cancer}}}{(2-s_{\text{host}})(2-s_{\text{cancer}}) + s_{\text{host}}\,s_{\text{cancer}}} \tag{22}$$

In other words, the equilibrium point (0.5,0.5) is **unstable if** $\alpha < A^*$

*Comparison between $A^*$ and $A_+$:* We can show that if the condition $\alpha < A^*$ is fulfilled, then $\alpha < A_+$:

$$\frac{A_+}{A^*} - 1 = \frac{U_1 + U_2}{2\,(2 - s_{\text{cancer}})\, s_{\text{host}}\, s_{\text{cancer}}} \tag{23}$$

With:

$$U_1 = s_{\text{host}}\,(1 - s_{\text{cancer}})\,(2 - s_{\text{host}})\,[(2 - s_{\text{host}})(2 - s_{\text{cancer}}) + s_{\text{host}}\, s_{\text{cancer}}] > 0 \tag{24}$$

$$\begin{aligned} U_2 = [(2 - s_{\text{host}})(2 - s_{\text{cancer}}) + s_{\text{host}}\, s_{\text{cancer}}] & \sqrt{s_{\text{host}}^2\,(1 - s_{\text{cancer}})^2\,(2 - s_{\text{host}})^2 + (2 - s_{\text{cancer}})\,(2 - s_{\text{host}})\, s_{\text{host}}\, s_{\text{cancer}}} \\ & - 2\,(2 - s_{\text{cancer}})\, s_{\text{host}}\, s_{\text{cancer}} \end{aligned} \tag{25}$$

We now determine the sign of $U_2$, which is the same as:

$$\begin{aligned} U_3 = [(2 - s_{\text{host}})(2 - s_{\text{cancer}}) + s_{\text{host}}\, s_{\text{cancer}}]^2 & \left[s_{\text{host}}^2\,(1 - s_{\text{cancer}})^2\,(2 - s_{\text{host}})^2 + (2 - s_{\text{cancer}})\,(2 - s_{\text{host}})\, s_{\text{host}}\, s_{\text{cancer}}\right] \\ & - 4\,(2 - s_{\text{cancer}})^2\, s_{\text{host}}^2\, s_{\text{cancer}}^2 = V_1 + V_2 \end{aligned} \tag{26}$$

With:

$$V_1 = [(2 - s_{\text{host}})(2 - s_{\text{cancer}}) + s_{\text{host}}\, s_{\text{cancer}}]^2 \left[s_{\text{host}}^2\,(1 - s_{\text{cancer}})^2\,(2 - s_{\text{host}})^2\right] > 0 \tag{27}$$

$$V_2 = [(2 - s_{\text{host}})(2 - s_{\text{cancer}}) + s_{\text{host}}\, s_{\text{cancer}}]^2\,[(2 - s_{\text{cancer}})\,(2 - s_{\text{host}})\, s_{\text{host}}\, s_{\text{cancer}}] - 4\,(2 - s_{\text{cancer}})^2\, s_{\text{host}}^2\, s_{\text{cancer}}^2 \tag{28}$$

We get:

$$V_2 = s_{\text{host}}\, s_{\text{cancer}}(2 - s_{\text{cancer}}) \left[4\,\left(2 - s_{\text{host}}\,(2 - s_{\text{host}})^2\right) s_{\text{cancer}}^2 - 8\,\left(4 - s_{\text{host}}\,\left(s_{\text{host}}^2 - 5\, s_{\text{host}} + 7\right)\right) s_{\text{cancer}} + 4\,(2 - s_{\text{cancer}})^3\right] \tag{29}$$

The polynomial function in the last term of this equation is a positive function for $0 < s_{\text{host}} < 1$ and $0 < s_{\text{cancer}} < 1$. Therefore:

$$V_2 > 0 \quad\longrightarrow\quad U_3 > 0 \quad\longrightarrow\quad U_2 > 0 \quad\longrightarrow\quad \frac{A_+}{A^*} - 1 > 0 \quad\longrightarrow\quad A^* < A_+ \tag{30}$$

The condition for unstability ($\alpha < A^*$) is true only if eigenvalues are complex. Here we showed that if the condition $\alpha < A^*$ is fulfilled, then the eigenvalues are necessarily complex ($\alpha < A_+$).

### APPENDIX B:
### Numerical analysis of a population genetic model with similarity selection

#### Purpose

The Red Queen Hypothesis for the evolution of sex traditionally relies on the existence of coevolutionary cycling. Nonetheless, parasite-host interactions can still promote the evolution of sex through a separate mechanism, known as similarity selection, that does not depend on genotypic selection such as would be imposed by coevolutionary fluctuations (Agrawal, 2006). Similarity selection occurs when there is a cost to being genotypically similar to one's family members. In particular, due to the transmission of parasites among family members, this cost exists if infection compatibility is under genetic influence (e.g., a matching alleles system). Therefore, vertical transmission of parasites leading to similarity selection can be a potent force favouring the evolution of sex and recombination (Agrawal, 2006).

We here modify the three-locus population genetic model of the manuscript to investigate whether similarity selection favours the evolution of sex (as in Agrawal, 2006) in the face of neoplasia dampening coevolutionary cycling.

#### Including vertical transmission and similarity selection

Just like in the three-locus population genetic model presented in the manuscript, we follow the genotypic frequencies of haploid hosts and haploid cancer cells through a life cycle that consists of a census, reproduction, neoplasia (development of neocancers), and selection.

During the reproduction phase, we now keep track of the association between offspring's and mother's genotypes. In particular, assuming that there is no mutation, offspring resulting from asexual reproduction always have the same genotype as their mother. By contrast, offspring resulting from sexual reproduction may have a different genotype as their mother.

During the selection phase, we keep track of the proportions of hosts of each genotype that have been infected by transmissible cancer (via genetic matching). This corresponds to the proportion of infected mothers of each genotype at the next selection phase.

During the selection phase, we now consider that a fraction $\Phi$ of hosts encounter transmissible cancers that successfully infected their mother (if any, those transmissible cancers therefore match the mother's genotype), while the other fraction $1 - \Phi$ encounter parasites at random (as in the main analysis). Parameter $\Phi$ therefore reflects the proportion of vertical transmission that causes similarity selection.

We fix the parameter $\Phi$, and we perform the same sensitivity analysis as in Figure 1.

#### Results

When we consider some extent of vertical transmission ($\Phi > 0$), a modifier allele associated with sexual reproduction can invade in an asexual population (as shown by Agrawal, 2006) (Fig. B1). We note that transmission among relatives does not dampen coevolutionary cycling as in Greenspoon and Mideo (2017); this is because we model explicitly vertical transmission by keeping track of the association between the mother-offspring genotypes.

Just like in the main analysis, even a small proportion of neocancers is sufficient to bring the coevolutionary dynamics between hosts and transmissible cancers to a halt. Nonetheless, similarity selection does not depend on coevolutionary fluctuations and favours sex (Fig. B1). Remember that we have not implemented any cost associated with sexual reproduction in this model, except the recombination load. Theoretical literature on similarity selection remains scarce, and whether similarity selection can overcome the two-fold cost of sex remains an open question. To our knowledge, three theoretical models on host-parasite interactions included similarity selection occurring via parental transmission of parasites (Agrawal, 2006; Greenspoon and M'Gonigle, 2013, 2014), and only one of those investigated its implication for sexual reproduction (Agrawal, 2006).

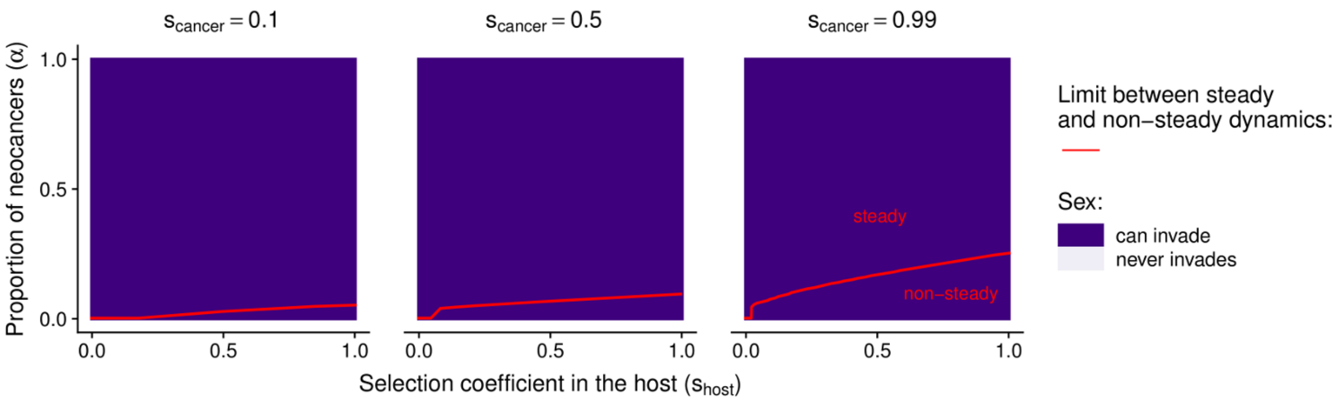


Figure B1: Coevolutionary dynamics between hosts and transmissible cancers, and evolution of sex when accounting for vertical transmission and similarity selection. Sensitivity of the population genetic models to the selection coefficients ($s_{\text{host}}, s_{\text{cancer}}$) and to the proportion of transmissible neocancers that are recently derived from the original host ($\alpha$). Red lines delimit the parameter spaces leading to non-steady and steady coevolutionary dynamics. The dynamic is defined as 'steady' when the variance in genotypic frequencies over 500 time steps is below $10^{-10}$. Dark purple indicates conditions under which a modifier allele associated with sexual reproduction (and with recombination, at least for one of the recombination rates tested) can invade in at least one of the 100 simulation runs. $\Phi = 0.1$.

### APPENDIX C:
### Analytical derivation of the epidemiological model

#### Purpose

We analyze the epidemiological model presented in the manuscript (Equation 4). Using basic linear algebra, we find the expression of densities at equilibrium and we show that this equilibrium state is locally stable. We also determine the expressions determining the proportion of neocancers ($\hat{\alpha}$) and the selection coefficient caused by infection by transmissible cancers ($\hat{s}_{\text{host}}$; inferred as the lifespan reduction in the host population due to infection by transmissible cancers) at equilibrium. Finally, we determine their sensitivity to changes in parameter values.

#### Differential equations

The following equations control the changes in densities of susceptible hosts ($S$), hosts that developed a neocancer by neoplasia ($I_0$), and hosts that are infected by a transmitted cancer ($I_{\text{T}}$):

$$\begin{cases} \dfrac{\mathrm{d}S}{\mathrm{d}t} = b\,(S + I_0 + I_{\text{T}})\left(1 - \dfrac{S + I_0 + I_{\text{T}}}{K}\right) - \left(\mu + \lambda_0 + \beta\,\dfrac{I_0 + I_{\text{T}}}{S + I_0 + I_{\text{T}}}\right) S \\ \dfrac{\mathrm{d}I_0}{\mathrm{d}t} = \lambda_0\;(S + \theta\, I_{\text{T}}) - \left(\mu + \nu + \theta\,\beta\,\dfrac{I_0 + I_{\text{T}}}{S + I_0 + I_{\text{T}}}\right) I_0 \\ \dfrac{\mathrm{d}I_{\text{T}}}{\mathrm{d}t} = \beta\,\dfrac{I_0 + I_{\text{T}}}{S + I_0 + I_{\text{T}}}\;(S + \theta\, I_0) - (\mu + \nu + \theta\,\lambda_0)\; I_T \end{cases} \tag{31}$$

With birth rate $b > 0$, carrying capacity $K > 0$, mortality rates $\mu > 0$ and $\nu > 0$, rate of neoplasia $\lambda_0 > 0$, transmission rate $\beta > 0$, and rate of changes in infection status $\theta \in [0, 1]$.

With the change of variable $N = S + I_0 + I_{\text{T}}$ and $I = I_0 + I_{\text{T}}$, the system of equations is equivalent to:

$$\begin{cases} \dfrac{\mathrm{d}N}{\mathrm{d}t} = b\,N\left(1 - \dfrac{N}{K}\right) - \mu\,N - \nu\,I \\ \dfrac{\mathrm{d}I}{\mathrm{d}t} = \left(\lambda_0 + \beta\,\dfrac{I}{N}\right)(N - I) - (\mu + \nu)\,I \\ \dfrac{\mathrm{d}I_0}{\mathrm{d}t} = \lambda_0\;(N - I + \theta\;(I - I_0)) - \left(\mu + \nu + \theta\,\beta\,\dfrac{I}{N}\right) I_0 \end{cases} \tag{32}$$

Below, we derive the equilibrium state of this system of equations.

#### Equilibrium

At equilibrium, we get:

$$\begin{cases} N^* = K\left(1 - \dfrac{\mu + P^*\nu}{b}\right) \\ I^* = P^*\,N^* \\ I_0^* = \dfrac{\lambda_0\,(1 - P^*)}{\mu + \nu} N^* \end{cases} \tag{33}$$

With $P^*$ the prevalence of transmissible cancers at equilibrium:

$$P^* = \frac{\beta - \lambda_0 - \mu - \nu + \sqrt{(\beta - \lambda_0 - \mu - \nu)^2 + 4\beta\lambda_0}}{2\,\beta} \in [0, 1] \tag{34}$$

To get $N^* > 0$, the condition $b > \mu + P^*\nu$ must be satisfied.
If $b \leq \mu + P^*\nu$, then $N^* \leq 0$ and the host population gets extinct.

***Calculation at equilibrium:***

Given that $\frac{\mathrm{d}N}{\mathrm{d}t} = 0$ at equilibrium, then $b\,N^* \left(1 - \frac{N^*}{K}\right) - \mu\,N^* - \nu\,I^* = 0$. Therefore:

$$I^* = \frac{1}{\nu} \left[ b \left(1 - \frac{N^*}{K}\right) - \mu \right] N^* \tag{35}$$

Given that $\frac{\mathrm{d}I}{\mathrm{d}t} = 0$ at equilibrium, then $\left(\lambda_0 + \beta\,\frac{I^*}{N^*}\right)(N^* - I^*) - (\mu + \nu)\,I^* = 0$, and therefore:

$$\beta\,{I^*}^2 - (\beta - \lambda_0 - \mu - \nu)\,N^* I^* - \lambda_0 {N^*}^2 = 0 \tag{36}$$

The only positive positive root of this polynomial equation is:

$$I^* = \frac{\beta - \lambda_0 - \mu - \nu + \sqrt{(\beta - \lambda_0 - \mu - \nu)^2 + 4\beta\lambda_0}}{2\,\beta}\,N^* = P^* N^* \tag{37}$$

This gives us the prevalence $P^*$ of transmissible cancers at equilibrium:

$$P^* = \frac{I^*}{N^*} = \frac{\beta - \lambda_0 - \mu - \nu + \sqrt{(\beta - \lambda_0 - \mu - \nu)^2 + 4\beta\lambda_0}}{2\,\beta} > 0 \tag{38}$$

$P^* < 1$ because:

$$1 - P^* = \frac{1}{2\,\beta} \left[ \beta + \lambda_0 + \mu + \nu - \sqrt{(\beta - \lambda_0 - \mu - \nu)^2 + 4\beta\lambda_0} \right] \tag{39}$$

and:

$$(\beta + \lambda_0 + \mu + \nu)^2 - \left((\beta - \lambda_0 - \mu - \nu)^2 + 4\beta\lambda_0\right) = 4\,\beta\,(\mu + \nu) > 0 \tag{40}$$

By combining equations 35 and 37, we get:

$$\frac{1}{\nu} \left[ b \left(1 - \frac{N^*}{K}\right) - \mu \right] N^* = P^* N^* \tag{41}$$

Which leads to the expression of $N^*$:

$$N^* = K \left(1 - \frac{\mu + P^* \nu}{b}\right) \tag{42}$$

And therefore to the expression of $I^*$:

$$I^* = P^* N^* = P^* K \left(1 - \frac{\mu + P^* \nu}{b}\right) \tag{43}$$

The host population does not get extinct if:

$$1 - \frac{\mu + P^* \nu}{b} > 0 \tag{44}$$

$$b > \mu + P^* \nu \tag{45}$$

Which is equivalent to the condition:

$$2\,\beta\,(b - \mu) - \nu\,(\beta - \lambda_0 - \mu - \nu) - \nu\,\sqrt{(\beta - \lambda_0 - \mu - \nu)^2 + 4\beta\lambda_0} > 0 \tag{46}$$

Given that $\frac{\mathrm{d}I_0}{\mathrm{d}t} = 0$ at equilibrium, then $\lambda_0\,(N^* - I^* + \theta\,(I^* - I_0^*)) - \left(\mu + \nu + \theta\,\beta\,\frac{I^*}{N^*}\right) I_0^* = 0$. With $I^* = P^* N^*$, we get:

$$I_0^* = \frac{\lambda_0\,(1 - P^* + \theta\,P^*)}{\mu + \nu + \theta\,(\lambda_0 + \beta P^*)}\,N^* \tag{47}$$

Which is equivalent to:

$$I_0^* = \frac{\lambda_0\,(1 - P^*)}{\mu + \nu} \times \frac{(\mu + \nu)\,(1 - P^* + \theta\,P^*)}{(1 - P^*)\,(\mu + \nu + \theta\,(\lambda_0 + \beta P^*))} \times N^* \tag{48}$$

$$I_0^* = \frac{\lambda_0\,(1-P^*)}{\mu+\nu} \times \frac{(1-P^*)\left(\mu+\nu+\theta\,\frac{(\mu+\nu)P^*}{1-P^*}\right)}{(1-P^*)\,(\mu+\nu+\theta\,(\lambda_0+\beta P^*))} \times N^* \tag{49}$$

Yet, $\frac{(\mu+\nu)P^*}{1-P^*} = \lambda_0 + \beta P^*$ because:

$$\frac{(1-P^*)\,(\lambda_0+\beta P^*)}{\mu+\nu} = \frac{\beta-\lambda_0-\mu-\nu+\sqrt{(\beta-\lambda_0-\mu-\nu)^2+4\beta\lambda_0}}{2\,\beta} = P^* \tag{50}$$

Therefore:

$$I_0^* = \frac{\lambda_0\,(1-P^*)}{\mu+\nu} N^* \tag{51}$$

### Local stability

Unlike in the population genetic model analyzed in Appendix A, we assume that time is continuous in the epidemiological model. At the equilibrium point, the local stability can therefore be inferred from the sign of the real part of the eigenvalues of the Jacobian matrix $\mathbf{J}$ at the equilibrium point $(N^*, I^*, I_0^*)$, the expression of which is:

$$\mathbf{J}_{\left(N^*,I^*,I_0^*\right)} = \begin{bmatrix} b-\mu-\frac{2\,b}{K}N^* & -\nu & 0 \\ \lambda_0+\beta\,P^{*2} & (\beta-\lambda_0-\mu-\nu)-2\,\beta\,P^* & 0 \\ \lambda_0+\theta\,\beta\,P^*\frac{\lambda_0(1-P^*)}{\mu+\nu} & \lambda_0\,(\theta-1)+\theta\,\beta\frac{\lambda_0(1-P^*)}{\mu+\nu} & -(\lambda_0\,\theta+\mu+\nu)-\theta\,\beta\,P^* \end{bmatrix} \tag{52}$$

Notably, the terms of the Jacobian matrix do not depend on $I^*$ and $I_0^*$. They depend however on $N^*$, and on $P^*$ the prevalence of transmissible cancers at equilibrium, calculated as:

$$P^* = \frac{\beta-\lambda_0-\mu-\nu+\sqrt{(\beta-\lambda_0-\mu-\nu)^2+4\beta\lambda_0}}{2\,\beta} \in [0,1] \tag{53}$$

The last column of the Jacobian matrix $\mathbf{J}_{\left(N^*,I^*,I_0^*\right)}$ is all zeros except for the element along the diagonal. This element, $-(\lambda_0\,\theta+\mu+\nu)-\theta\,\beta\,P^*$, is therefore one of the eigenvalues. This eigenvalue is negative. The remaining eigenvalues are the eigenvalues of the smaller matrix:

$$\mathbf{J'}_{\left(N^*,I^*,I_0^*\right)} = \begin{bmatrix} b-\mu-\frac{2\,b}{K}N^* & -\nu \\ \lambda_0+\beta\,P^{*2} & (\beta-\lambda_0-\mu-\nu)-2\,\beta\,P^* \end{bmatrix} \tag{54}$$

We can infer the sign of real parts of the remaining eigenvalues from the signs of the determinant and the trace of the matrix. In particular, the real parts of the remaining eigenvalues are both negative (which is the condition for equilibria to be stable) if: $\det\left(\mathbf{J'}_{\left(\mathbf{N^*},\mathbf{I^*},\mathbf{I_0^*}\right)}\right) > 0$ and $\operatorname{trace}\left(\mathbf{J'}_{\left(\mathbf{N^*},\mathbf{I^*},\mathbf{I_0^*}\right)}\right) < 0$.

At equilibrium, $N^* = K\left(1-\frac{\mu+P^*\nu}{b}\right)$, and we can calculate:

$$\det\left(\mathbf{J'}_{\left(N^*,I^*,I_0^*\right)}\right) = \frac{\sqrt{(\beta-\lambda_0-\mu-\nu)^2+4\beta\lambda_0}}{2\,\beta}\left[2\,\beta\,(b-\mu)-\nu\,(\beta-\lambda_0-\mu-\nu)-\nu\,\sqrt{(\beta-\lambda_0-\mu-\nu)^2+4\beta\lambda_0}\right] \tag{55}$$

Following the condition of existence of the equilibrium (Equation 46), the second term of this expression is positive, and therefore:

$$\det\left(\mathbf{J'}_{\left(N^*,I^*,I_0^*\right)}\right) > 0 \tag{56}$$

We also get the expression of the trace of the matrix:

$$\operatorname{trace}\left(\mathbf{J'}_{\left(N^*,I^*,I_0^*\right)}\right) = -b+\beta-\lambda_0+\nu+2\,(\nu-\beta)\,P^* \tag{57}$$

$$\begin{aligned}\operatorname{trace}\left(\mathbf{J'}_{\left(N^*,I^*,I_0^*\right)}\right) = \frac{1}{\beta}\Bigg[&-\left(2\,\beta\,(b-\mu)-\nu\,(\beta-\lambda_0-\mu-\nu)-\nu\,\sqrt{(\beta-\lambda_0-\mu-\nu)^2+4\beta\lambda_0}\right) \\ &+\beta\left(b-\mu-\sqrt{(\beta-\lambda_0-\mu-\nu)^2+4\beta\lambda_0}\right)\Bigg]\end{aligned} \tag{58}$$

If $b - \mu - \sqrt{(\beta - \lambda_0 - \mu - \nu)^2 + 4\beta\lambda_0} \leq 0$, we can conclude from Equation 58 that the trace of the matrix $\mathbf{J'}_{\left(\mathbf{N^*},\mathbf{I^*},\mathbf{I_0^*}\right)}$ is negative (given that the condition of existence of the equilibrium if fulfilled; Equation 46).

The trace can also be expressed as:

$$\text{trace}\left(\mathbf{J'}_{\left(N^*,I^*,I_0^*\right)}\right) = -\left(b - \mu - \sqrt{(\beta - \lambda_0 - \mu - \nu)^2 + 4\beta\lambda_0}\right) + 2\left(\nu\, P^* - \sqrt{(\beta - \lambda_0 - \mu - \nu)^2 + 4\beta\lambda_0}\right) \tag{59}$$

As detailed below, we can show that $\nu\, P^* - \sqrt{(\beta - \lambda_0 - \mu - \nu)^2 + 4\beta\lambda_0} < 0$. Therefore, the trace is also negative if $b - \mu - \sqrt{(\beta - \lambda_0 - \mu - \nu)^2 + 4\beta\lambda_0} > 0$. Thus, the trace of the matrix $\mathbf{J'}_{\left(\mathbf{N^*},\mathbf{I^*},\mathbf{I_0^*}\right)}$ is always negative under the condition of existence of the equilibrium.

Given that $\det\left(\mathbf{J'}_{\left(\mathbf{N^*},\mathbf{I^*},\mathbf{I_0^*}\right)}\right) > 0$ and $\text{trace}\left(\mathbf{J'}_{\left(\mathbf{N^*},\mathbf{I^*},\mathbf{I_0^*}\right)}\right) < 0$ under the condition of existence of the equilibrium, the real parts of the two remaining eigenvalues of the Jacobian matrix at equilibrium are both negative (just like the first eigenvalue; determined directly from the expression of the Jacobian matrix, Equation 52). **The equilibrium point is therefore locally stable**.

***Proof that*** $\nu\, P^* - \sqrt{(\beta - \lambda_0 - \mu - \nu)^2 + 4\beta\lambda_0} < 0$ ***:***

Here we use the notation $X = (\beta - \lambda_0 - \mu - \nu)^2 + 4\beta\lambda_0$, and we aim at showing that $\nu\, P^* - \sqrt{X} < 0$.
$\nu\, P^* - \sqrt{X}$ has the same sign as: $\nu^2\, {P^*}^2 - X$. We can calculate:

$$\nu^2\, {P^*}^2 - X = \frac{1}{4\,\beta^2}\left[(\nu - 2\beta)\,(\nu + 2\beta)\,\sqrt{X}^2 + 2\nu^2\,(\beta - \lambda_0 - \mu - \nu)\,\sqrt{X} + \nu^2\,(\beta - \lambda_0 - \mu - \nu)^2\right] \tag{60}$$

We call $Q$ the polynomial function:

$$Q(x) = (\nu - 2\beta)\,(\nu + 2\beta)\,x^2 + 2\nu^2\,(\beta - \lambda_0 - \mu - \nu)\,x + \nu^2\,(\beta - \lambda_0 - \mu - \nu)^2 \tag{61}$$

Therefore, $\nu\, P^* - \sqrt{X}$ has the same sign as the polynomial fonction $Q$ at $x = \sqrt{X}$.
The discriminant of this polynomial function $Q$ is positive:

$$\Delta_Q = 16\nu^2\beta^2\,(\beta - \lambda_0 - \mu - \nu)^2 > 0 \tag{62}$$

And there are therefore two values $x_1$ and $x_2$ that are solution to the equation $Q(x) = 0$:

$$x_1 = \frac{-\nu\,(\beta - \lambda_0 - \mu - \nu)}{\nu + 2\beta} \tag{63}$$

$$x_2 = \frac{-\nu\,(\beta - \lambda_0 - \mu - \nu)}{\nu - 2\beta} \tag{64}$$

If $\nu - 2\beta \geq 0$, then we have necessarily $\beta - \lambda_0 - \mu - \nu \leq 0$, and $x_1 > 0$, $x_2 > 0$, hence $x_1 < x_2$. In that condition, $Q(x) < 0$ if $x \in [x_1, x_2]$.
For $\nu - 2\beta \geq 0$, we get:

$$x_1^2 - \sqrt{X}^2 = \frac{-4\beta\left[(\nu + \beta)\,(\beta - \lambda_0 - \mu - \nu)^2 + \lambda_0\right]}{(\nu + 2\beta)^2} < 0 \tag{65}$$

$$x_2^2 - \sqrt{X}^2 = \frac{4\beta\left[(\nu - 2\beta)\left[(\nu - 2\beta)\,(\mu + \nu - \beta) - (\beta - \lambda_0 - \mu - \nu)\,(\beta + \lambda_0 + \mu)\right] + \beta\,(\beta - \lambda_0 - \mu - \nu)^2\right]}{(\nu - 2\beta)^2} > 0 \tag{66}$$

This means that if $\nu - 2\beta \geq 0$, we get $\sqrt{X} \in [x_1, x_2]$ and $Q(\sqrt{X}) < 0$. Therefore, in that condition, we get $\nu\, P^* - \sqrt{X} < 0$.

If $\nu - 2\beta < 0$, then $x_1$ and $x_2$ have opposite signs, and for any $x > 0$, we get $Q(x) < 0$ if $x > \max(x_1, x_2)$

For $\beta - \lambda_0 - \mu - \nu \geq 0$, we get $x_1 < 0$ and $x_2 > 0$, hence $\max(x_1, x_2) = x_2$. Yet, $\sqrt{X} > x_2$ because, in that case, we have:

$$x_2^2 - \sqrt{X}^2 = \frac{4\beta \left[ (\nu - \beta)(\beta - \lambda_0 - \mu - \nu)^2 - \lambda_0 (\nu - 2\beta)^2 \right]}{(\nu - 2\beta)^2} < 0 \tag{67}$$

Indeed, for $\beta - \lambda_0 - \mu - \nu > 0$, we have necessarily $\nu - \beta < 0$.
For $\beta - \lambda_0 - \mu - \nu < 0$, we get $x_1 > 0$ and $x_2 < 0$, hence $\max(x_1, x_2) = x_1$. Yet, $\sqrt{X} > x_1$ because, in that case, we have:

$$x_1^2 - \sqrt{X}^2 = \frac{-4\beta \left[ (\nu + \beta)(\beta - \lambda_0 - \mu - \nu)^2 + \lambda_0 \right]}{(\nu + 2\beta)^2} < 0 \tag{68}$$

This means that if $\nu - 2\beta < 0$, we get $\sqrt{X} > \max(x_1, x_2)$ and $Q(\sqrt{X}) < 0$. Therefore, in that condition, we get again $\nu P^* - \sqrt{X} < 0$.

### Proportion of neocancers at equilibrium

At equilibrium, we calculate the proportion of neocancers as:

$$\hat{\alpha} = \frac{I_0^*}{I^*}. \tag{69}$$

Therefore:

$$\boxed{\hat{\alpha} = \frac{\lambda_0}{\mu + \nu} \left( \frac{1}{P^*} - 1 \right)} \tag{70}$$

### Selection coefficient due to transmissible cancers at equilibrium

At equilibrium, we infer the selection coefficient as the lifespan reduction due to the risk of being infected by transmissible cancers:

$$\boxed{\hat{s}_{\text{host}} = \frac{1}{1 + \frac{\mu}{\nu} \frac{1}{P^*}}} \tag{71}$$

***Calculation:***
The mean time spent in a each state ($S$, $I_0$, $I_{\text{T}}$, cf. Equation 31) is:

$$t_S = \frac{1}{\mu + \lambda_0 + \beta \frac{I^*}{N^*}} \tag{72}$$

$$t_{I_0} = \frac{1}{\mu + \nu + \theta\, \beta \frac{I^*}{N^*}} \tag{73}$$

$$t_{I_{\text{T}}} = \frac{1}{\mu + \nu + \theta\, \lambda_0} \tag{74}$$

As a newly infected individual (by either a neocancer or a transmitted cancer), you live:

$$\begin{cases} T_{I_0} = t_{I_0} + \dfrac{\theta\, \beta \frac{I^*}{N^*}}{\mu + \nu + \theta\, \beta \frac{I^*}{N^*}}\, t_{I_{\text{T}}} \\ T_{I_{\text{T}}} = t_{I_{\text{T}}} + \dfrac{\theta\, \lambda_0}{\mu + \nu + \theta\, \lambda_0}\, t_{I_0} \end{cases} \tag{75}$$

Which gives us:

$$T_{I_0} = T_{I_{\text{T}}} = \frac{1}{\mu + \nu} \tag{76}$$

As a new born, you live:

$$T_S = t_S + \frac{\lambda_0}{\mu + \lambda_0 + \beta \frac{I^*}{N^*}} T_{I_0} + \frac{\beta \frac{I^*}{N^*}}{\mu + \lambda_0 + \beta \frac{I^*}{N^*}} T_{I_\mathrm{T}} \tag{77}$$

$$T_S = \frac{1}{\mu + \lambda_0 + \beta \frac{I^*}{N^*}} \left(1 + \frac{\lambda_0 + \beta \frac{I^*}{N^*}}{\mu + \nu}\right) \tag{78}$$

Without transmissible cancers, an individuals has an average lifespan equal to $1/\mu$. The lifespan reduction due to the risk of being infected by transmissible cancers is therefore:

$$\hat{s}_\mathrm{host} = 1 - \frac{T_S}{1/\mu} \tag{79}$$

$$\hat{s}_\mathrm{host} = \frac{\nu \left(\lambda_0 + \beta \frac{I^*}{N^*}\right)}{(\mu + \nu)\left(\mu + \lambda_0 + \beta \frac{I^*}{N^*}\right)} \tag{80}$$

Yet, given that $\frac{\mathrm{d}I}{\mathrm{d}t} = 0$ at equilibrium, then $\left(\lambda_0 + \beta \frac{I^*}{N^*}\right)(N^* - I^*) - (\mu + \nu) I^* = 0$, and $\beta \frac{I^*}{N^*} = \frac{(\mu+\nu)I^*}{N^* - I^*} - \lambda_0$ and therefore:

$$\hat{s}_\mathrm{host} = \frac{\nu I^*}{\mu N^* + \nu I^*} \tag{81}$$

This means that the lifespan reduction due to the risk of being infected by transmissible cancers is equivalent to the relative mortality rate caused by cancer in the population.

Finally, given that $I^* = P^* N^*$ at equilibrium:

$$\hat{s}_\mathrm{host} = \frac{1}{1 + \frac{\mu}{\nu}\frac{1}{P^*}} \tag{82}$$

### Effect of b, K and $\theta$

While parameters $b$ and $K$ change the densities $N^*$, $I^*$, ${I_0}^*$ at equilibrium, they do not affect $P^*$, $\hat{\alpha}$ and $\hat{s}_\mathrm{host}$.
Parameter $\theta$ has no effect on the densities $N^*$, $I^*$, ${I_0}^*$ at equilibrium, and does not affect $P^*$, $\hat{\alpha}$ and $\hat{s}_\mathrm{host}$.

### Effect of $\nu$

***On the prevalence of transmissible cancers:***
We determine the effect of $\nu$ on the prevalence of transmissible cancers:

$$\frac{\partial P^*}{\partial \nu} = \frac{-P^*}{\sqrt{(\beta - \lambda_0 - \mu - \nu)^2 + 4\beta\lambda_0}} < 0 \tag{83}$$

An increase in the cancer-associated mortality rate increases the mortality rate of infected hosts, thereby decreasing the prevalence of transmissible cancers at equilibrium.

***On the proportion of neocancers:***
We determine the effect of $\nu$ on the proportion of neocancers at equilibrium:

$$\frac{\partial \hat{\alpha}}{\partial \nu} = \lambda_0 \left[\frac{\partial \frac{1}{\mu+\nu}}{\partial \nu}\left(\frac{1}{P^*} - 1\right) + \frac{1}{\mu + \nu}\frac{\partial\left(\frac{1}{P^*} - 1\right)}{\partial \nu}\right] \tag{84}$$

The cancer-associated mortality rate decreases both the densities of hosts infected by a neocancers ($I_0$) and hosts infected by transmitted cancer ($I_\mathrm{T}$) at equilibrium. As shown by the signs of the terms $\frac{\partial \frac{1}{\mu+\nu}}{\partial \nu} < 0$ and $\frac{\partial\left(\frac{1}{P^*} - 1\right)}{\partial \nu} > 0$, the cancer-associated mortality rate affects the proportion of neocancer $\hat{\alpha}$ depending on those effects on $I_0$ and $I_0 + I_\mathrm{T}$.

$$\frac{\partial \hat{\alpha}}{\partial \nu} = \frac{\lambda_0 \left[\mu + \nu - (1 - P^*) \sqrt{(\beta - \lambda_0 - \mu - \nu)^2 + 4\beta\lambda_0}\right]}{(\mu + \nu)^2 P^* \sqrt{(\beta - \lambda_0 - \mu - \nu)^2 + 4\beta\lambda_0}} \tag{85}$$

$$\frac{\partial \hat{\alpha}}{\partial \nu} = \frac{\lambda_0 \left[(\beta - \lambda_0 - \mu - \nu)^2 + 2\,\beta\,(\mu + \nu + 2\,\lambda_0) - (\beta + \lambda_0 + \mu + \nu) \sqrt{(\beta - \lambda_0 - \mu - \nu)^2 + 4\beta\lambda_0}\right]}{2\,\beta\,(\mu + \nu)^2 P^* \sqrt{(\beta - \lambda_0 - \mu - \nu)^2 + 4\beta\lambda_0}} \tag{86}$$

Yet,

$$\left[(\beta - \lambda_0 - \mu - \nu)^2 + 2\,\beta\,(\mu + \nu + 2\,\lambda_0)\right]^2 - (\beta + \lambda_0 + \mu + \nu)^2 \left((\beta - \lambda_0 - \mu - \nu)^2 + 4\beta\lambda_0\right) = 4\,\beta^2(\mu + \nu)^2 > 0 \tag{87}$$

And therefore:

$$\frac{\partial \hat{\alpha}}{\partial \nu} > 0 \tag{88}$$

A high cancer-associated mortality rate decreases the density of hosts infected by transmitted cancers relatively more than the density of hosts infected by neocancers. This makes sense because a high mortality rate of infected hosts decreases the density of hosts infected by transmitted cancers via both increased mortality and reduced transmission. Overall, increased cancer-associated mortality rate increases the proportion of neocancers.

***On the selection coefficient:***
The cancer-associated mortality rate has antagonistic effects on the selection coefficient due to transmissible cancers:

$$\frac{\partial \hat{s}_{\text{host}}}{\partial \nu} = \frac{\mu}{(P^*\,\nu + \mu)^2} \frac{\partial\,(\nu P^*)}{\partial \nu} \tag{89}$$

As emphasized by the term $\frac{\partial(\nu P^*)}{\partial \nu}$, a high cancer-associated mortality rate increases mortality of infected hosts, but also decreases the cancer prevalence (which increases $\hat{s}_{\text{host}}$). The balance between those antagonistic effects will determine whether increased cancer-associated mortality rate decreases or increases the selection coefficient due to transmissible cancers – i.e, whether $\frac{\partial \hat{s}_{\text{host}}}{\partial \nu} < 0$ or $> 0$.

$$\frac{\partial \hat{s}_{\text{host}}}{\partial \nu} = \frac{\mu\,P^*}{(P^*\,\nu + \mu)^2} \times \left(1 - \frac{\nu}{\sqrt{(\beta - \lambda_0 - \mu - \nu)^2 + 4\beta\lambda_0}}\right) \tag{90}$$

Therefore, $\frac{\partial \hat{s}_{\text{host}}}{\partial \nu}$ has the same sign as:

$$\left((\beta - \lambda_0 - \mu - \nu)^2 + 4\beta\lambda_0\right) - \nu^2 = 2\,(\mu + \lambda_0 - \beta)\,(\mu + \nu) + (\lambda_0 + \beta)^2 - \mu^2, \tag{91}$$

and $\frac{\partial \hat{s}_{\text{host}}}{\partial \nu} > 0$ for:

$$\begin{cases} \mu + \nu \leq \dfrac{\mu^2 - (\lambda_0 + \beta)^2}{2\,(\mu + \lambda_0 - \beta)}, & \text{for} \quad \mu \leq \beta - \lambda_0 \\ \mu + \nu > \dfrac{\mu^2 - (\lambda_0 + \beta)^2}{2\,(\mu + \lambda_0 - \beta)}, & \text{for} \quad \mu > \beta - \lambda_0 \end{cases} \tag{92}$$

Yet, because $\mu + \nu > \mu$, we can show that:

$$\mu > \beta - \lambda_0 \implies \mu + \nu > \frac{\mu^2 - (\lambda_0 + \beta)^2}{2\,(\mu + \lambda_0 - \beta)} \tag{93}$$

And therefore, $\frac{\partial \hat{s}_{\text{host}}}{\partial \nu} > 0$ for :

$$\begin{cases} \mu \leq \mu + \nu \leq \dfrac{\mu^2 - (\lambda_0 + \beta)^2}{2\,(\mu + \lambda_0 - \beta)}, & \text{for} \quad \mu \leq \beta - \lambda_0 \\ \mu + \nu \geq \mu, & \text{for} \quad \mu > \beta - \lambda_0 \end{cases} \tag{94}$$

$$\begin{cases} 0 \leq \nu \leq \dfrac{(\beta - \mu - \lambda_0)^2 + 4\,\lambda_0\,\beta}{2(\beta - \mu - \lambda_0)}, & \text{for} \;\; \mu \leq \beta - \lambda_0 \\ \qquad\qquad\qquad\qquad \nu \geq 0, & \text{for} \;\; \mu > \beta - \lambda_0 \end{cases} \tag{95}$$

Increased cancer-associated mortality rate increases the selection coefficient as long as the mortality rate of infected individuals ($\mu+\nu$) is low enough. Otherwise the reduction of prevalence decreases the selection coefficient due to transmissible cancers.

#### Effect of $\mu$

Given that the expressions of $P^*$ and $\hat{\alpha}$ depends on $\mu+\nu$. The parameter $\mu$ has the same effect on $P^*$ and $\hat{\alpha}$ as parameter $\nu$ by affecting the mortality rate of infected individuals.

***On the prevalence of transmissible cancers:***

$$\frac{\partial P^*}{\partial \mu} = \frac{\partial P^*}{\partial \nu} < 0 \tag{96}$$

An increase in the baseline mortality rate increases the mortality rate of infected hosts, thereby decreasing the prevalence of transmissible cancers at equilibrium.

***On the proportion of neocancers:***

$$\frac{\partial \hat{\alpha}}{\partial \mu} = \frac{\partial \hat{\alpha}}{\partial \nu} > 0 \tag{97}$$

An increase in the baseline mortality rate increases the mortality rate of infected hosts, thereby increasing the proportion of neocancers.

***On the selection coefficient:***
Increased baseline mortality rate decreases the selection coefficient due to cancer by reducing both the prevalence of transmissible cancers ($P^*$) and the relative mortality cost associated with cancer ($\nu/\mu$).

$$\frac{\partial \hat{s}_{\text{host}}}{\partial \mu} = \frac{-\nu\,P^*}{\left(P^*\,\nu + \mu\right)^2}\left[1 + \frac{\mu}{\sqrt{\left(\beta - \lambda_0 - \mu - \nu\right)^2 + 4\beta\lambda_0}}\right] < 0 \tag{98}$$

#### Effect of $\lambda_0$

***On the prevalence of transmissible cancers:***
We determine the effect of $\lambda_0$ on the prevalence of transmissible cancers:

$$\frac{\partial P^*}{\partial \lambda_0} = \frac{1}{2\,\beta\,\sqrt{\left(\beta - \lambda_0 - \mu - \nu\right)^2 + 4\beta\lambda_0}}\left[\beta + \lambda_0 + \mu + \nu - \sqrt{\left(\beta - \lambda_0 - \mu - \nu\right)^2 + 4\beta\lambda_0}\right] \tag{99}$$

Yet,

$$\left(\beta + \lambda_0 + \mu + \nu\right)^2 - \left(\left(\beta - \lambda_0 - \mu - \nu\right)^2 + 4\beta\lambda_0\right) = 4\,\beta\,(\mu + \nu) > 0 \tag{100}$$

Therefore:

$$\frac{\partial P^*}{\partial \lambda_0} > 0 \tag{101}$$

The rate of neoplasia increases the prevalence of transmissible cancers at equilibrium.

***On the proportion of neocancers:***
We determine the effect of $\lambda_0$ on the proportion of neocancers at equilibrium:

$$\frac{\partial \hat{\alpha}}{\partial \lambda_0} = \frac{1}{\mu+\nu}\left[\left(\frac{1}{P^*}-1\right) + \lambda_0 \frac{\partial\left(\frac{1}{P^*}-1\right)}{\partial \lambda_0}\right] \tag{102}$$

The rate of neoplasia increases both the densities of hosts infected by a neocancers ($I_0$) and hosts infected by transmitted cancer ($I_{\mathrm{T}}$) at equilibrium. As shown by the sign of the terms $\frac{1}{P^*}-1>0$ and $\lambda_0 \frac{\partial\left(\frac{1}{P^*}-1\right)}{\partial \lambda_0}<0$, the rate of neoplasia affects the proportion of neocancer $\hat{\alpha}$ depending on those effects on $I_0$ and $I_0+I_{\mathrm{T}}$.

$$\frac{\partial \hat{\alpha}}{\partial \lambda_0} = \frac{1-P^*}{2\,\beta\,(\mu+\nu)\,{P^*}^2\sqrt{(\beta-\lambda_0-\mu-\nu)^2+4\beta\lambda_0}}\left[(\beta-\lambda_0-\mu-\nu)^2+2\,\beta\,\lambda_0+(\beta-\lambda_0-\mu-\nu)\sqrt{(\beta-\lambda_0-\mu-\nu)^2+4\beta\lambda_0}\right] \tag{103}$$

Yet,

$$\left[(\beta-\lambda_0-\mu-\nu)^2+2\,\beta\,\lambda_0\right]^2-(\beta-\lambda_0-\mu-\nu)^2\left((\beta-\lambda_0-\mu-\nu)^2+4\beta\lambda_0\right)=4\,\beta^2\,{\lambda_0}^2>0 \tag{104}$$

And therefore:

$$\frac{\partial \hat{\alpha}}{\partial \lambda_0}>0 \tag{105}$$

A high rate of neoplasia increases the density of hosts infected by neocancers relatively more than the density of hosts infected by transmitted cancers. Overall, increased rate of neoplasia increases the proportion of neocancers.

***On the selection coefficient:***
A high rate of neoplasia increases the selection coefficient due to transmissible cancers by increasing the prevalence:

$$\frac{\partial \hat{s}_{\mathrm{host}}}{\partial \lambda_0} = \frac{\mu}{\nu\left(1+\frac{\mu}{\nu}\frac{1}{P^*}\right)^2 {P^*}^2}\frac{\partial P^*}{\partial \lambda_0}>0 \tag{106}$$

#### Effect of $\beta$

***On the prevalence of transmissible cancers:***
We determine the effect of $\beta$ on the prevalence of transmissible cancers:

$$\frac{\partial P^*}{\partial \beta} = \frac{1}{2\,\beta^2\sqrt{(\beta-\lambda_0-\mu-\nu)^2+4\beta\lambda_0}}\left[(\lambda_0+\mu+\nu)\sqrt{(\beta-\lambda_0-\mu-\nu)^2+4\beta\lambda_0}-\left((\lambda_0+\mu+\nu)^2+\beta\,(\lambda_0-\mu-\nu)\right)\right] \tag{107}$$

Yet,

$$(\lambda_0+\mu+\nu)^2\left((\beta-\lambda_0-\mu-\nu)^2+4\beta\lambda_0\right)-\left((\lambda_0+\mu+\nu)^2+\beta\,(\lambda_0-\mu-\nu)\right)^2=4\,\beta^2\lambda_0\,(\mu+\nu)>0 \tag{108}$$

Therefore:

$$\frac{\partial P^*}{\partial \beta}>0 \tag{109}$$

The transmission rate increases the prevalence of transmissible cancers at equilibrium.

***On the proportion of neocancers:***
We determine the effect of $\beta$ on the proportion of neocancers at equilibrium:

$$\frac{\partial \hat{\alpha}}{\partial \beta} = -\frac{\lambda_0}{(\mu+\nu){P^*}^2}\frac{\partial P^*}{\partial \beta}<0 \tag{110}$$

A high transmission rate increases the density of hosts infected by transmitted cancers. Therefore, increased transmission rate decreases the proportion of neocancers.

***On the selection coefficient:***

A high transmission rate increases the selection coefficient due to transmissible cancers by increasing the prevalence:

$$\frac{\partial \hat{s}_{\text{host}}}{\partial \beta} = \frac{\mu}{\nu \left(1 + \frac{\mu}{\nu} \frac{1}{P^*}\right)^2 P^{*2}} \frac{\partial P^*}{\partial \beta} > 0 \tag{111}$$

### APPENDIX D:
### Numerical analysis of an epidemiological model with two types of cancers

#### Purpose

We modify the epidemiological model from the manuscript (Equation 4) to account for distinct types of transmissible cancers differing genetically (such as in the population genetic model we analyzed). Here we consider that there are two types of transmissible cancers that originate from different types of hosts (e.g., with different genetic makeup). We assume that hosts are susceptible only to the cancerous line that is originally derived from an host of their own type (because this cancerous line has the same genetic make up than their own and is not recognized as non-self).

We show below that this model has the same behaviour as the epidemiological model analyzed in the main text. In particular, we show that there is no coevolutionary cycling in this epidemiological model where time is assumed to be continuous (unlike in our population genetic model). We also highlight that by ignoring diversity in cancerous strains, we underestimated the proportion of neocancers and we overestimated the strength of selection caused by transmissible cancers. The epidemiological constraints caused by the coexistence of multiple cancerous lines therefore reduce even further the conditions under which transmissible cancers can promote the evolution of sex.

#### Differential equations

The following equations control the changes in densities of those two types of hosts. Again, we follow the densities of susceptible hosts ($S_1$ and $S_2$), hosts that developed a neocancer by neoplasia ($I_{01}$ and $I_{02}$), and hosts that are infected by a transmitted cancer ($I_{\mathrm{T}1}$ and $I_{\mathrm{T}2}$):

$$\begin{cases} \dfrac{\mathrm{d}S_1}{\mathrm{d}t} = b\,(S_1 + I_{01} + I_{\mathrm{T}1})\left(1 - \dfrac{N}{K}\right) - \left(\mu + \lambda_0 + \beta\,\dfrac{I_{01} + I_{\mathrm{T}1}}{N}\right) S_1 \\ \dfrac{\mathrm{d}I_{01}}{\mathrm{d}t} = \lambda_0\,(S_1 + \theta\, I_{\mathrm{T}1}) - \left(\mu + \nu + \theta\,\beta\,\dfrac{I_{01} + I_{\mathrm{T}1}}{N}\right) I_{01} \\ \dfrac{\mathrm{d}I_{\mathrm{T}1}}{\mathrm{d}t} = \beta\,\dfrac{I_{01} + I_{\mathrm{T}1}}{N}\,(S_1 + \theta\, I_{01}) - (\mu + \nu + \theta\,\lambda_0)\, I_T \\ \dfrac{\mathrm{d}S_2}{\mathrm{d}t} = b\,(S_2 + I_{02} + I_{\mathrm{T}2})\left(1 - \dfrac{N}{K}\right) - \left(\mu + \lambda_0 + \beta\,\dfrac{I_{02} + I_{\mathrm{T}2}}{N}\right) S_2 \\ \dfrac{\mathrm{d}I_{02}}{\mathrm{d}t} = \lambda_0\,(S_2 + \theta\, I_{\mathrm{T}2}) - \left(\mu + \nu + \theta\,\beta\,\dfrac{I_{02} + I_{\mathrm{T}2}}{N}\right) I_{02} \\ \dfrac{\mathrm{d}I_{\mathrm{T}2}}{\mathrm{d}t} = \beta\,\dfrac{I_{02} + I_{\mathrm{T}2}}{N}\,(S_2 + \theta\, I_{02}) - (\mu + \nu + \theta\,\lambda_0)\, I_T \end{cases} \tag{112}$$

With $N = S_1 + I_{01} + I_{\mathrm{T}1} + S_2 + I_{02} + I_{\mathrm{T}2}$.

With birth rate $b > 0$, carrying capacity $K > 0$, mortality rates $\mu > 0$ and $\nu > 0$, rate of neoplasia $\lambda_0 > 0$, transmission rate $\beta > 0$, and rate of changes in infection status $\theta \in [0, 1]$.

#### Numerical analysis

We could not analyze the above model with analytical derivations. Instead, we analyzed it numerically.

We implemented initial differences in prevalence between the two cancer strains by drawing initial densities ($S_1$, $S_2$, $I_{01}$, $I_{02}$, $I_{\mathrm{T}1}$, $I_{\mathrm{T}2}$)) from a uniform distribution across the range [1, $K/6$]. Using Julia (version 1.0.1), we then ran simulations until time $t = 5,000$.

We tested the same parameter values as in Figure 4 in the manuscript (with $b = 1.0$, $K = 500$, $\theta = 0$). For each combinations of parameters, we ran two replicates characterized by different initial densities to investigate the existence of an unstable equilibrium state.

At the end of the simulation, we calculated the prevalence as:

$$\text{Prevalence} = \frac{I_{01} + I_{02} + I_{\text{T}1} + I_{\text{T}2}}{N} \tag{113}$$

We also calculated the proportion of neocancers ($\hat{\alpha}$) as:

$$\hat{\alpha} = \frac{I_{01} + I_{02}}{I_{01} + I_{02} + I_{\text{T}1} + I_{\text{T}2}} \tag{114}$$

Finally, we assessed the selection coefficient caused by transmissible cancers ($\hat{s}_{\text{host}}$) as the relative mortality rate caused by cancer in the population (cf. Equation 81 in Appendix C):

$$\hat{s}_{\text{host}} = \frac{(I_{01} + I_{02} + I_{\text{T}1} + I_{\text{T}2})\,\nu}{(I_{01} + I_{02} + I_{\text{T}1} + I_{\text{T}2})\,(\mu + \nu) + (S_1 + S_2)\,\mu} \tag{115}$$

### Results

***Nature of the equilibrium state:***

If the system was characterized by an unstable equilibrium, we would observe variations among the replicate runs that differed in their initial state (either because of cycling or chaotic behaviours, or because of convergence towards distinct stable equilibria). Here, however, we get the same $\hat{\alpha}$ and $\hat{s}_{\text{host}}$ values at the end of the two replicate ran for each combination of parameters (Fig. D1). Therefore, we can conclude that the system is characterized by only one stable equilibrium, just like the simpler system we analyzed in the main text. Overlapping generations and epidemiological dynamics (with changes in parasite prevalence) are probably the causes behind this dampening of coevolutionary cycling among hosts and cancer strains (as shown in other models; e.g., May and Anderson, 1983; Beck, 1984; Kouyos et al., 2007; MacPherson and Otto, 2018, cited in our manuscript).

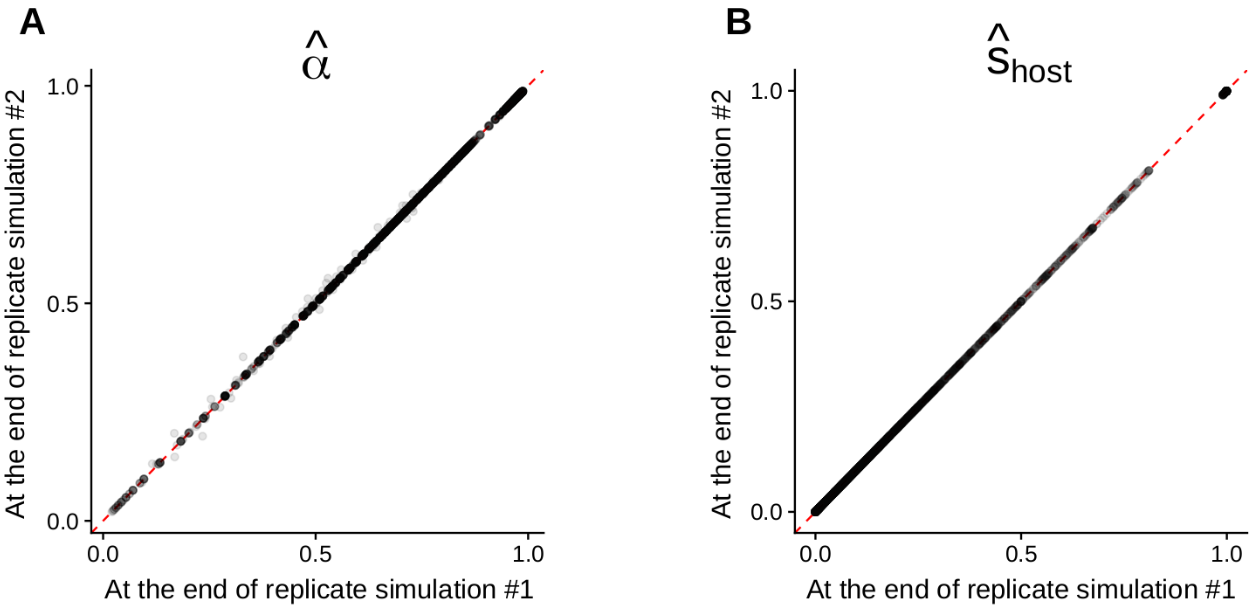


Figure D1: Comparison of the output ($\hat{\alpha}$ and $\hat{s}_{\text{host}}$ values at the end of the simulations) of two replicates differing by their initial conditions. We implement point transparency to ignore errors due to the numerical approximation inherent to the differential equations solver. Parameter values as in Fig. D2.

***Effects of epidemiological parameters:***

Epidemiological parameters have similar effects on the proportion of neocancers ($\hat{\alpha}$) and the selection coefficient caused by transmissible cancers ($\hat{s}_{\text{host}}$) at equilibrium than in the original model with only one type of transmissible cancer (Fig. D2 vs. Fig. 4A-B). Nonetheless, the prevalence of cancer is particularly low when we account for two types of transmissible cancers (Fig. D3A) because the transmission rate is lower (given that hosts can avoid infection by half of the cancerous lines). This is particularly the case when the rate of neoplasia ($\lambda_0$) is low (Fig. D3A). This leads to a high proportion of neocancers (Fig. D3B) and a low selection coefficient caused by transmissible cancers (Fig. D3C) compared to the situation with only one type of transmissible cancers. This indirect effect on $\hat{\alpha}$ and $\hat{s}_{\text{host}}$ driven by changes in prevalence was highlighted in the original model (cf. Fig. 3).

Therefore, the epidemiological constraints caused by the coexistence of multiple cancerous lines constrain even further the conditions under which transmissible cancers can promote the evolution of sex.

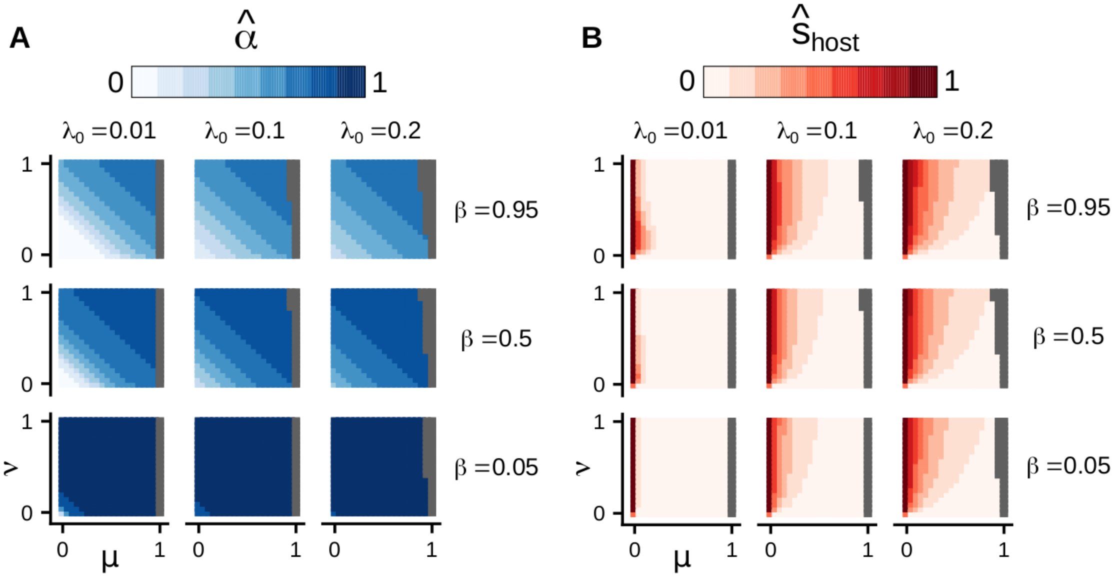


Figure D2: Effects of epidemiological parameters on $(\hat{\alpha}, \hat{s}_{\text{host}})$ at the end of the simulation when we account for two types of transmissible cancers. We represent in gray the conditions under which $N < 1$ at the end of the simulation – i.e., we can thus assume that the host population gets extinct for these combinations of parameters. Parameter values: $b = 1.0$, $K = 500$, $\theta = 0$. Note that we get the exact same results when we implement other values of parameters $b$, $K$ or $\theta$ (not shown).

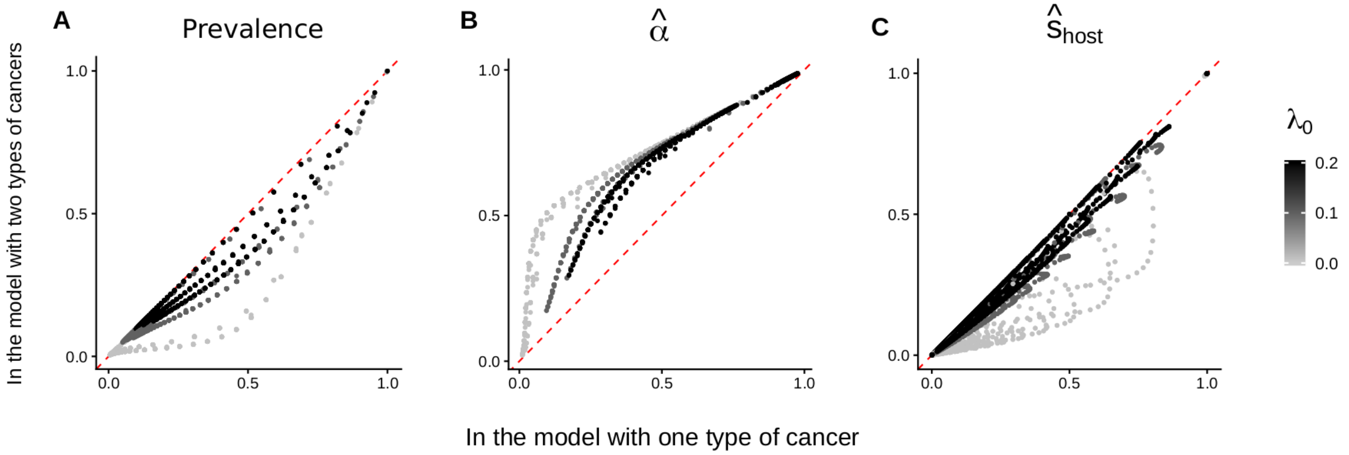


Figure D3: Comparison with the situation with only one type of cancer. We compare the output of this model with the values from the model where we considered only type of cancer. The gray scale represents the value of $\lambda_0$ (rate of neoplasia) implemented. Parameter values as in Fig. D2.

### References


Agrawal A. F., 2006. Similarity selection and the evolution of sex: Revisiting the red queen. *PLoS Biology*, 4(8):1364–1371. doi: 10.1371/journal.pbio.0040265.

Beck K., 1984. Coevolution: mathematical analysis of host-parasite interactions. *Journal of Mathematical Biology*, 19(1):63–77. doi: 10.1007/BF00275931.

Greenspoon P. B. and M'Gonigle L. K., 2013. The evolution of mutation rate in an antagonistic coevolutionary model with maternal transmission of parasites. *Proceedings of the Royal Society B: Biological Sciences*, 280(1761):20130647. doi: 10.1098/rspb.2013.0647.

Greenspoon P. B. and M'Gonigle L. K., 2014. Host-parasite interactions and the evolution of nonrandom mating. *Evolution*, 68(12):3570–3580. doi: 10.1111/evo.12538.

Greenspoon P. B. and Mideo N., 2017. Parasite transmission among relatives halts Red Queen dynamics. *Evolution*, 71(3):747–755. doi: 10.1111/evo.13157.

Hickey D. A. and Golding G. B., 2018. The advantage of recombination when selection is acting at many genetic Loci. *Journal of Theoretical Biology*, 442:123–128. doi: 10.1016/j.jtbi.2018.01.018.

Iles M. M., Walters K., and Cannings C., 2003. Recombination can evolve in large finite populations given selection on sufficient loci. *Genetics*, 165(4):2249–2258.

Kouyos R. D., Salathé M., and Bonhoeffer S., 2007. The Red Queen and the persistence of linkage-disequilibrium oscillations in finite and infinite populations. *BMC Evolutionary Biology*, 7:211. doi: 10.1186/1471-2148-7-211.

MacPherson A. and Otto S. P., 2018. Joint coevolutionary-epidemiological models dampen Red Queen cycles and alter conditions for epidemics. *Theoretical Population Biology*, 122:137–148. doi: 10.1016/j.tpb.2017.12.003.

May R. M. and Anderson R. M., 1983. Epidemiology and genetics in the coevolution of parasites and hosts. *Proceedings of the Royal Society B: Biological Sciences*, 219(1216):281–313. doi: 10.1098/rspb.1983.0075.

Otto S. P. and Nuismer S. L., 2004. Species interactions and the evolution of sex. *Science*, 304(5673):1018–1020. doi: 10.1126/science.1094072.
